# Attenuation of Motion Artifacts in fMRI using Discrete Reconstruction of Irregular fMRI Trajectories (DRIFT)

**DOI:** 10.1101/714436

**Authors:** David Parker, Qolamreza Razlighi

## Abstract

Numerous studies reported motion as the most detrimental source of noise and artifacts in functional magnetic resonance imaging (fMRI). Different approaches have been proposed and used to attenuate the effect of motion on fMRI data, including both prospective and retrospective (post-processing) techniques. However, each type of motion (e.g. translation versus rotation or in-plane versus out-of-plane) has a distinct effect on the MR signal, which is not fully understood nor appropriately modeled in the field. In addition, effects of the same motion can be substantially different depending on when it occurs during the pulse sequence (e.g. RF excitation, gradient encoding, or k-space read-out). Thus, each distinct kind of motion and the time of its occurrence may require a unique approach to be optimally corrected. Therefore, we start with an investigation of the effects of different motions on the MR signal based on the Bloch equation. We then simulate their unique effects with a comprehensive fMRI simulator. Our results indicate that current motion correction methods fail to completely address the motion problem. Retrospective techniques such as spatial realignment can correct for between-volume misalignment, but fail to address within volume contamination and spin-history artifacts. Because of the steady state nature of the fMRI acquisition, spin-history artifacts arising from over/under excitation during slice-selection causes the motion artifacts to contaminate MR signal even after cessation of motion, which makes it challenging to be corrected retrospectively. Prospective motion correction has been proposed to prevent spin-history artifacts, but fails to address motion artifacts during k-space readout. In this article, we propose a novel method to remove these artifacts: Discrete reconstruction of irregular fMRI trajectory (DRIFT). Our method calculates the exact displacement of k-space recording due to motion at each dwell time and retrospectively corrects each slice of the fMRI volume using an inverse nonuniform Fourier transform. We evaluate our proposed methods using simulated data as well as fMRI data collected from a rotating phantom inside a 3T Siemens Prisma scanner. We conclude that a hybrid approach with both prospective and retrospective components are essentially required for optimal removal of motion artifacts from the fMRI data.

## Introduction

Motion is one of the most prominent sources of noise and artifacts in fMRI. Significant amounts of work has been devoted to reduce the artifacts of motion in the fMRI signal (Hedley, Yan and Rosenfeld, 1991; Kim *et al.*, 1999; Oakes *et al.*, 2005; Johnstone *et al.*, 2006; Bhagalia and Kim, 2008; White *et al.*, 2010; Power *et al.*, 2014; Zaitsev, Maclaren and Herbst, 2015; Chunli *et al.*, 2015; Birn, 2016; Zahneisen *et al.*, 2016). Presently, there are two major directions of research in the fMRI motion problem: One direction is to remove motion related variance from motion contaminated data (retrospective correction), while the other is to prevent motion artifacts from occurring in the first place (prospective correction).

Perhaps the most common retrospective correction technique used in the field is the correction of spatial misalignment. Spatial misalignment is a consequence of head movement inside the scanner from volume to volume. Misalignment causes voxels to sample different locations in the brain over time. Spatial realignment using rigid body registration is utilized to correct for misalignment retrospectively as a preprocessing step in many fMRI processing pipelines (Friston *et al.*, 1996). While rigid-body registration operates based on the assumption that head motion only occurs between volumes, in reality, motion continuously contaminates entirely/partially the acquired slices, and/or volumes, and may cause artifacts that persist after motion has ceased. Therefore, the effectiveness of spatial realignment is limited only to the volumes acquired well before and after the occurrence of motion. Though some slice-based registration methods exist (Kim *et al.*, 1999; Bannister, Michael Brady and Jenkinson, 2007), they still will be unable to recover signal from regions that are under-sampled due to motion (such as part of the head moving out of the field of view (FOV) of the scan).

Typically, spatial realignment is unable to remove all motion related variance from the fMRI timeseries. Because of this, it is also common for researchers to attempt to remove the remaining motion-induced variance from the data by regressing out motion parameters from fMRI timeseries. In theory, this should orthogonalize (remove any correlation between) the fMRI timeseries and the motion parameters. A major challenge with orthogonalization is that the relationship between motion-related changes in MR signal and motion parameters are extremely non-linear and complex. Thus, often times researchers also orthogonalize fMRI timeseries with time-delayed, derivatives and/or squared motion parameters (Friston *et al.*, 1996; Power *et al.*, 2014), averaged signals from CSF or white matter voxels (Jo *et al.*, 2010, 2013), or motion-related independent component analysis (ICA) components’ timeseries (Griffanti *et al.*, 2014). Additionally, advanced multi-echo pulse sequences have been used which allow researchers to separate BOLD (T_2_*) signal fluctuations from other sources of signal fluctuation, including those due to motion. By separating these signals, a more accurate regressor describing motion variance can be used to orthogonalize the fMRI data (Kundu *et al.*, 2012, 2014). Despite all these advancements, one of the most effective existing retrospective techniques involves censoring all the volumes acquired not only during motion but also before and after (Power *et al.*, 2014).

Prospective techniques (motion artifact prevention) aim to prevent these sources of variance from ever contaminating the fMRI signal. These techniques are desirable because they not only prevent misalignment, but also prevent another, even more destructive motion artifact known as “spin-history” artifacts (Friston *et al.*, 1996; Power *et al.*, 2014). Spin-history artifacts arise due to the fact that fMRI operates in a steady-state by periodically exciting a slice with a relatively short repetition time (TR) which does not allow a full recovery of the net magnetization. If the steady-state of a system is perturbed, it can have a lasting effect on many volumes into the future, until steady-state can be reached again. Any over/under excitation due to motion-related displacement of the slice will disrupt the equilibrium of the steady-state acquisition, which causes spin-history artifacts. This can be caused with out-of-plane translation or rotation. These motions are particularly harmful, as they introduce both within-volume misalignments that distort the reconstructed image as well as spin-history artifacts that alter the magnitude of the signal. An example of spin-history artifacts are shown in Figure 1. The top row shows the effect of out-of-plane rotation that occurs half-way through an acquisition, while the bottom row shows out-of-plane translation. The first column shows an image sampled without motion, where each slice is represented by a different color. The center two columns show the effect of motion on the true location of the sampled slices, as well as the resulting reconstructed image. Note that both spin-history artifacts *and* slice-misalignment are present in this reconstruction, and that this kind of intra-volume misalignment results in a distorted image that cannot be corrected for with rigid body registration. The final column shows the volume immediately after motion. In this volume, spin-history artifacts are still preset (both over and under-excitation), and the volume is misaligned, however the reconstruction is not distorted, and so it can be corrected with rigid body registration. However, retrospective correction for spin-history artifacts is extremely challenging, and to our knowledge, there is no existing method to effectively restore the underlying signal once contaminated with spin-history artifacts (Friston *et al.*, 1996; Yancey *et al.*, 2011; Schulz *et al.*, 2014; Zaitsev, Maclaren and Herbst, 2015). Prospective methods (which prevent spin-history artifacts) address this issue. In prospective motion correction (PMC), the scanner coordinate system (defined by the gradient orientation) is constantly updated to “follow” any motion the subject’s head may undergo. The optimal prospective method would require real-time motion tracking and gradient adjustment at each dwell time, which poses a technical challenge to be implemented with the current technology. However, current motion tracking and gradient adjustment are available for slice-by-slice application, and are implemented in several research laboratories (Thesen *et al.*, 2000; Zaitsev *et al.*, 2004, 2017; Speck, Hennig and Zaitsev, 2006; White *et al.*, 2010; Tisdall *et al.*, 2012; Schulz *et al.*, 2014; Todd *et al.*, 2015; Maclaren *et al.*, 2018; Frost *et al.*, 2019).

**Figure 1:**
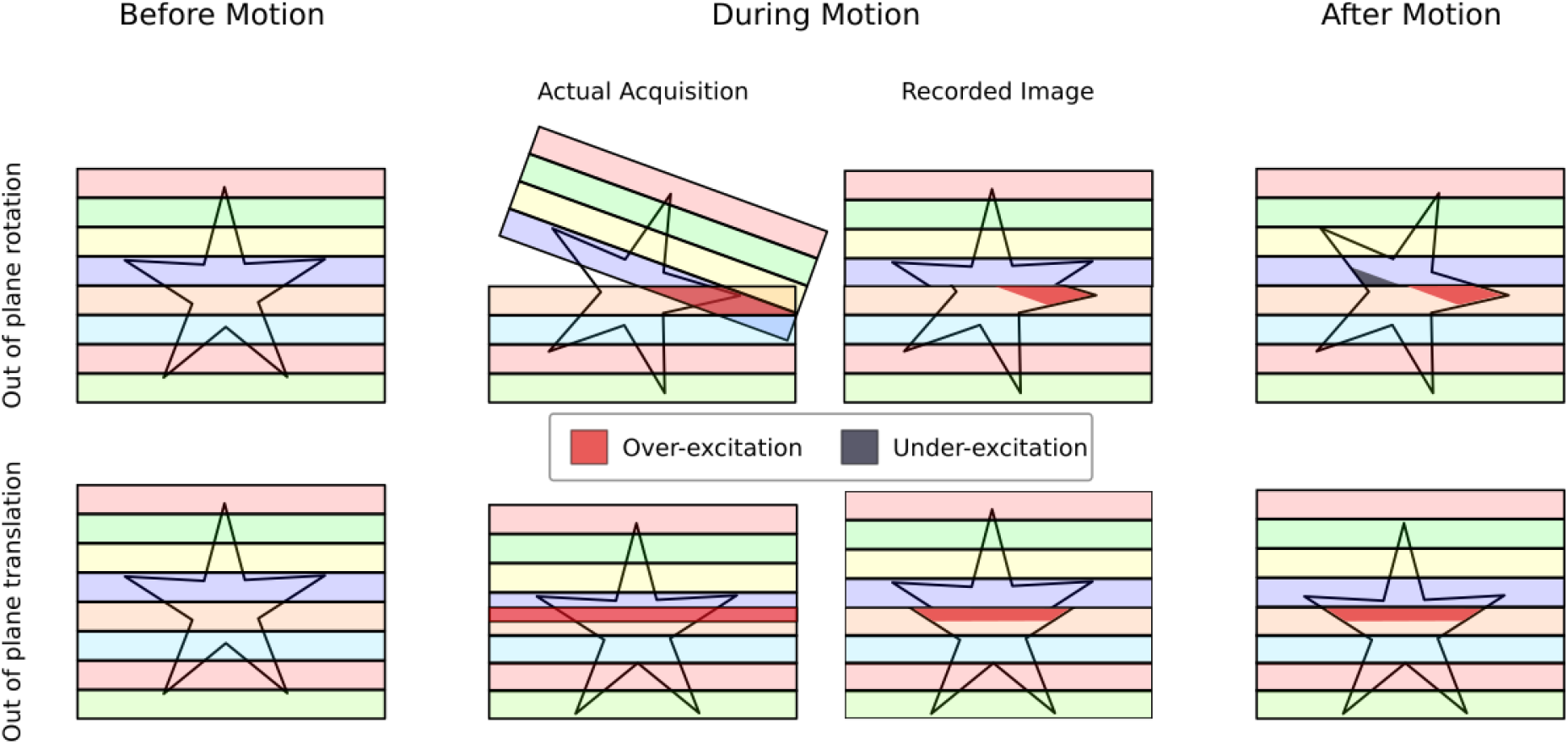
An illustration of spin-history artifacts for out-of-plane rotation and out-of-plane translation. An object is sampled slice by slice, from top to bottom, where each slice is shown as a colored bar. Without motion, the entire object is sampled normally without any artifacts, as shown in the first column. The center two columns represent image acquisition with motion, where the motion occurs half way through the acquisition of the object. The “Actual Acquisition” shows the actual location of the sampled slices for the given motion, while the “Recorded Image” shows the resulting reconstruction of the motion-contaminated volume. Finally, the column on the right shows the acquisition of the next volume, which is motionless, but is misaligned and has spin-history artifacts.

In its current state, slice-based prospective techniques leave an important motion artifact uncorrected: the motion induced disruption of k-space filling. K-space filling artifacts occur when motion disrupts the uniform sampling of k-space. Artifacts such as blurring, ghosting, or ringing in the reconstructed image are common effects of k-space displacement. Because of the small magnitude of the BOLD signal relative to the motion artifacts, it is imperative that these artifacts be optimally corrected or removed. Therefore, these artifacts will still need to be addressed even if spin-history artifacts are prevented. Given the current state of PMC, it is clear that a hybrid method is necessary to create an artifact free time series: a prospective correction of the gradients to prevent spin-history artifacts, and a retrospective correction to remove any remaining artifacts due to distorted k-space filling.

In this article, we first show that each kind of motion (such as translation versus rotation or in-plane versus out-of-plane) has a distinct effect on the acquired image based on the theory of MR signal generation and the Bloch equation. Some of the findings were striking, for instance out-of-plane translation is shown to have zero effect on the MR signal during k-space filling. Using the results of our analysis, we propose a retrospective motion correction technique: Discrete reconstruction of irregular fMRI trajectory (DRIFT), to correct for within-slice motion artifacts. DRIFT does not require a real-time implementation. Instead our method retrospectively estimates the motion-related k-space displacement and places each k-space sample at its true location according to the type and amount of motion observed at each dwell time. In the presence of motion, the new k-space sampling points may not fall on a regular and uniformly spaced grid which prevents use of the inverse fast Fourier transform (IFFT) to reconstruct the images. Instead, we used an irregular inverse Fourier transform (IIFT) that is irregular in the frequency domain, but discrete in the spatial domain. An IIFT is significantly more time consuming, but it can be applied retrospectively and without a need for real-time implementation. To show the validity of our findings, and to demonstrate the effectiveness of our method, we used a comprehensive fMRI simulator to generate artifacts from various motion profiles. We evaluated the effectiveness of our proposed method by correcting every different type of motion, individually and in combination, using our fMRI simulator. We have also evaluated our method using a rotating phantom inside 3T Siemens Prisma scanner.

## Materials & Methods

### Theory

To study the effect of motion on the MR signal, it is essential to independently investigate the effect of each type of motion using the theory of MR signal generation and the MR signal equation. By examining the different components present in standard MR signal equation, the effect of motion can be modeled mathematically, and the signal change can be evaluated for any specific kind of motion. The signal detected by the coils in the MRI scanner is dependent on the proton density ρ of all the tissue excited by the latest RF pulse at a spatial location ****r****. This signal is commonly approximated as:

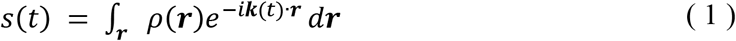

where ***k***(*t*) is the k-space coordinate, defined as:

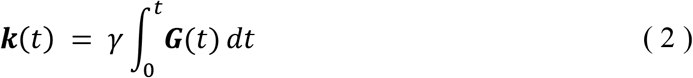

where *γ* is the gyromagnetic ratio and ***G***(*t*) is the gradient vector, describing the strength of the magnetic gradients along the *x*, *y*, and *z* axes at time *t*. This equation demonstrates how a sequence of applied gradients change the location in the k-space over time. The pattern in which consecutive points of k-space are sampled during EPI is called the k-space “trajectory”, and the time between two consecutive sampling points in k-space is called the dwell time. The k-space trajectory traces a path along k-space, sampling the MR signal at designated points along the way. When each designated point in the trajectory has been acquired, a complete image can be reconstructed. This happens over a very short period of time, typically 60ms, which is twice the echo time (TE) of the fMRI pulse sequence.

The most common trajectory for sampling k-space in fMRI is a Cartesian grid. A Cartesian grid trajectory can be seen in Figure 2 (a), with its reconstructed image in (b). If the grid has uniformly spaced samples, the image can be reconstructed with an IFFT. Without loss of generality, we will assume for the rest of this article that the *x* axis is the frequency encoding direction, *y* is the phase encoding direction, and *z* is the slice selection direction.

**Figure 2:**
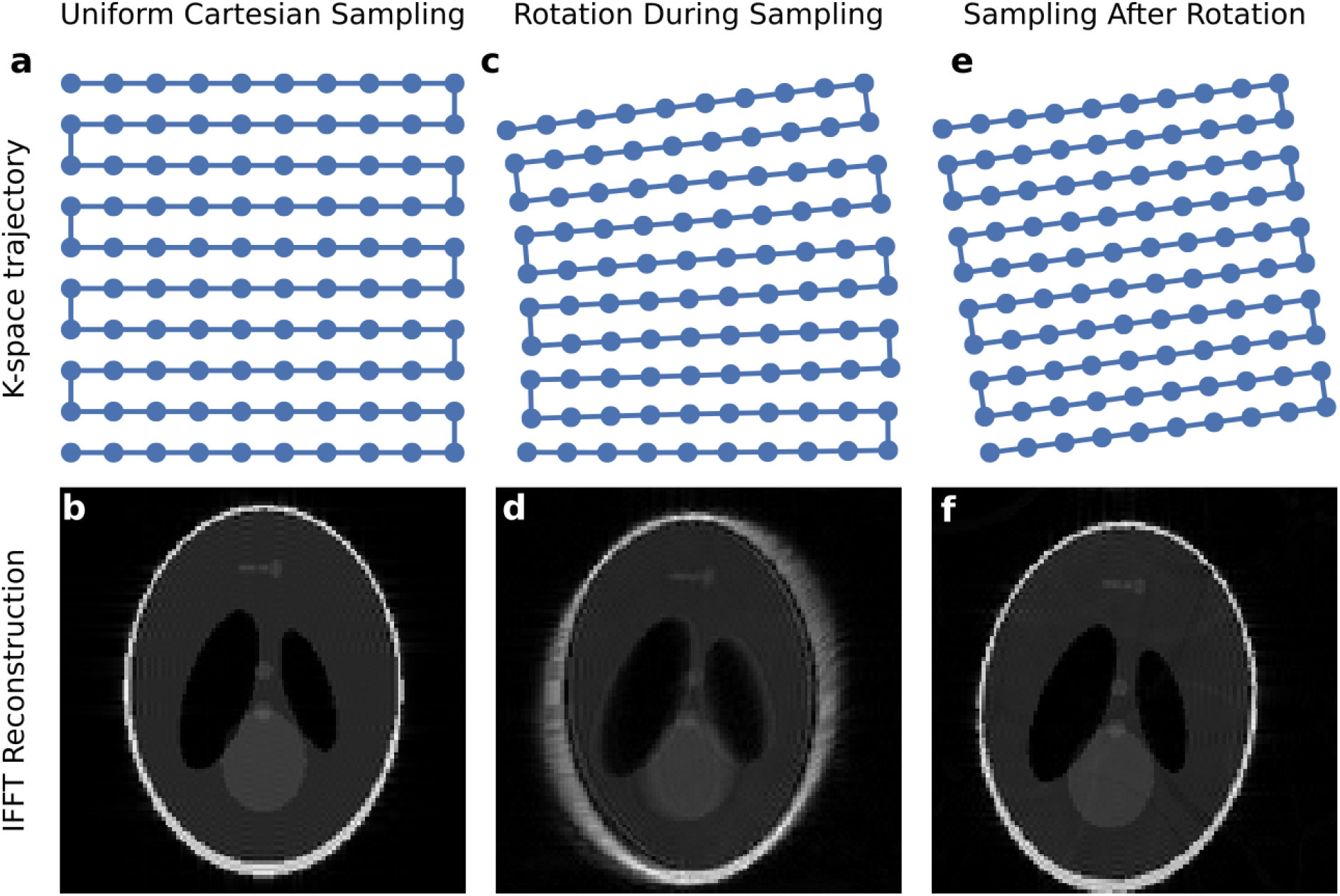
An illustration of the effect of motion on k-space sampling, and their associated artifacts. (A) The EPI pulse sequence is designed to sample an evenly-spaced Cartesian grid in k-space (B) which results in a clean reconstruction. (C) Motion *during* K-space readout (in this instance, in-plane rotation) results in a distorted k-space. (D) This leads to artifacts in the reconstructed image. (E) K-space readout of a stationary, but rotated image is equivalent to sampling a rotated but uniform grid. (F) This leads to a reconstructed image without artifacts, but that is rotated by the inverse of the k-space rotation.

### Formulating the Effect of Motion on the MR signal

We can now consider two coordinate systems: The object, ***r***_*ob*_, and the scanner, ***r***_*sc*_. Without motion, these coordinate systems are static, and we simply set them equal to each other, where ***r*** = ***r***_*ob*_ = ***r***_*sc*_. However, this no longer holds true if the object moves. When motion is present, we must describe the object’s location relative to its initial position at the time of excitation from the most recent RF pulse. If the object is initially aligned with the scanner’s coordinate system, ***r***_*sc*_, and moves in a way described by a rotation matrix R and a translation vector ***T***, we can describe a point in the object’s coordinate system in terms of the scanner’s coordinates:

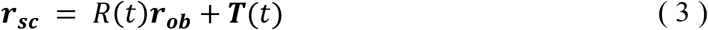

and it follows that we can describe a point in the scanner in terms of the object’s coordinates:

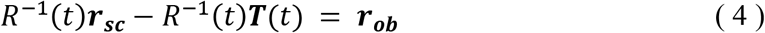

This means a rotation of the object in the scanner is equivalent to the scanner making an inverse rotation while the object is stationary (Drobnjak *et al.*, 2006). Treating the motion in the latter way actually results in a simplified analysis. To model the motion as a stationary object with an inverse rotation of the scanner, we re-derive the k-space equation to account for this rotation, as the spatial coordinate ***r*** is now changing with time. This process, along with the original derivation of k-space, is outlined in appendix A, and the results are shown here:

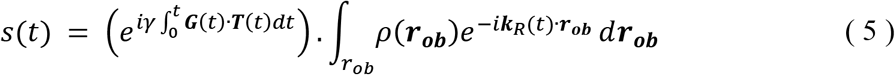

where ***k***_*R*_(*t*) is the new k-space location, which includes the motion related displacement, and is defined as:

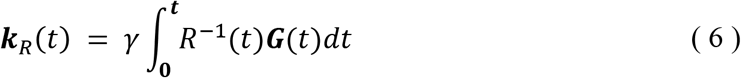

There is also an additional translation term which appears in front of the integral as it has no dependence on the spatial location ***r***. Equations (5) and (6) can now be used to examine the effects of different types of motion on the MR signal during k-space filling.

### Effect of Translational Movement on MR signal

We first begin by examining the exponential term outside the integral, which corresponds to the effect of translation. To understand the motion artifact due to translation from a theoretical point of view, we can expand the dot products in equation (5), and reformulate the exponential in terms of its gradient vector components:

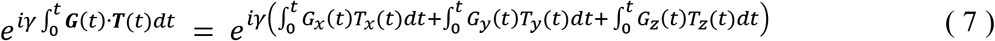

where *G*_*x*_(*t*) and *T*_*x*_(*t*) are the components of the gradient and translation vectors along the *x* axis, respectively. The same notation is also used for the gradient and translation along the *y* and *z* axes. As seen in equation (7), a gradient must be present along a given axis for translation in that direction to have any effect on the MR signal. However, a typical EPI pulse sequence uses only the frequency and phase encoding gradients during readout, and the slice selection gradient *G*_*z*_(*t*) is off during the entire period of k-space filling, making any translation along the *z* axis irrelevant to the MR signal. Artifacts due to translation along *z* only occur when the next z gradient is applied, which is usually concurrent with the next RF pulse, causing spin-history artifacts. Thus, if the subject’s motion can be tracked, and the gradients can be adjusted to compensate for motion before the next RF pulse, the out-of-plane translation will have negligible effect on the acquired MR signal. This is not surprising since every tissue in the slice would still experience the same *x* and *y* gradient magnitude as it moves along the slice selection axis. On the other hand, translation along the *x* and *y* axes will cause a proportional phase shift in the acquired MR signal. This can be corrected for by multiplying the detected signal by an inverse phase shift to cancel out the artifacts of the translational motion. For bulk motion, the entire k-space is multiplied by a single phase shift, while for gradual motion occurring during k-space filling, each point in k-space is multiplied with a unique phase shift, dependent on the subject’s motion at that point in time.

### Effect of Rotational Movement on MR signal

We now examine the effects of in-plane rotation *θ*_*z*_ around the *z* axis, and out-of-plane rotation *θ*_*x*_, and *θ*_*y*_ around the *x*, and *y* axes, respectively. For simplicity of notation, we drop the time index in this calculation and separately formulate the effect of each rotation by multiplying the gradient vector ***G*** by the respective rotation matrices for a rotation around each axis:

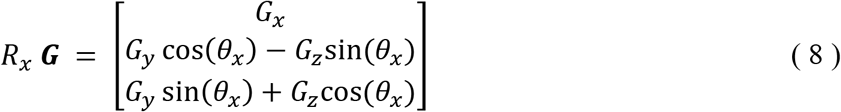

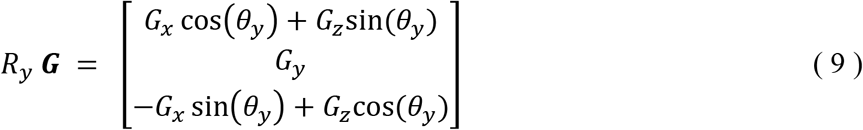

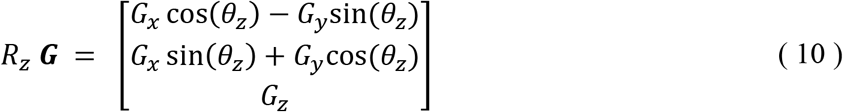

Again, since there is no active gradient along *z* axis during the k-space filling, we can further simplify the above equation for rotational motion.

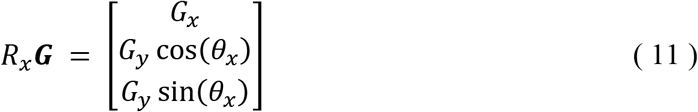

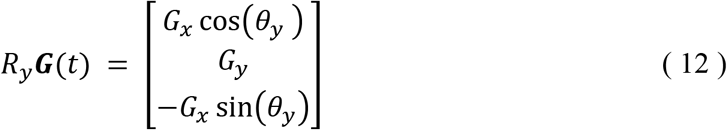

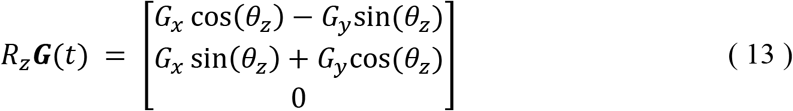

These equations demonstrate that both in-plane and out-of-plane rotations will introduce displacement to the sampled k-space locations. In the case of Cartesian sampling, the samples no longer lie on the expected uniformly spaced grid. This is illustrated in Figure 2 (c), which demonstrates an exaggerated in-plane rotation that occurs during the k-space acquisition. Figure 2 (d) shows the reconstructed image of this rotated k-space, with the associated reconstruction artifact for in-plane rotation. In the next excitation of this slice, if the gradients are not prospectively adjusted, the k-space samples will be uniform, but rotated, as shown in Figure 2 (e). The consequence of this is misalignment of the reconstructed image, shown in Figure 2 (f). It is important to note that the in-plane rotation artifact is limited to the slice in which the motion occurs, and will not contaminate future slices, other than misalignment.

The absence of a *z* gradient significantly reduces the magnitude of k-space sampling artifacts caused by an out-of-plane rotation around either the *x* or *y* axis. Equation (11) demonstrates that for typical rotations during a single k-space filling (usually less than 2 degrees) the magnitude of the gradient experienced by the object will remain around 99% of the original gradients, with only a small fraction appearing in the object’s *z* axis, since cos(2°) = 0.999, and sin(2°) = 0.035. Further, the gradient experienced along the axis of rotation remains unchanged, meaning if the object rotates around the *x* axis, it still experiences the same *x* gradient as it did before rotation. However, rotation around the *z* axis changes both the *x* and *y* gradients experienced by the object, and creates larger k-space sampling artifacts. In other words, we expect that in-plane rotation to generate larger displacement in the k-space locations, and therefore larger artifacts, than the out-of-plane rotation. To summarize the major theoretical points made in this section: 1) Translation along an axis can only cause signal artifacts if a gradient is present along the same axis. 2) Translation can be corrected for by multiplying the raw data with the appropriate phase shifts. 3) Rotation can be modeled directly as a change in k-space coordinates, and 4) In-plane rotation is significantly more detrimental than out-of-plane rotation.

### Discrete Reconstruction of Irregular fMRI Trajectory (DRIFT)

We propose to use the derived equations for motion contaminated k-space to modify the classical fMRI reconstruction algorithm in a way that accounts for and removes all motion artifacts. Presently, all fMRI images are reconstructed using an IFFT, shown in equation (14).

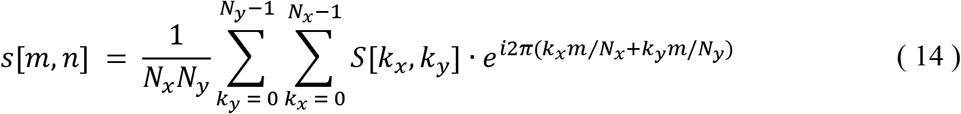

Where *s* is the reconstructed image at spatial coordinate [m, n], *S*[*k*_*x*_, *k*_*y*_] is the sampled k-space point at k-coordinate *k*_*x*_, *k*_*y*_, and N_x_, N_y_ is the total number of samples in the *x* and *y* axis respectively. In the IFFT, m, n, *k*_*x*_, and *k*_*y*_ are discrete values, and it is assumed that every sample is evenly spaced in the range k = 0 to N-1, hence the relative frequency contribution of each sample is k/N of the maximum frequency.

While the IFFT algorithm fails to reconstruct the motion contaminated image, we can modify equation (14) to account for samples that are offset from their customary k/N location. If the motion is accurately measured, the true location of each k-space sample can be calculated from equation (6). It is at this point that most reconstruction steps would “regrid”, by interpolating k-space to estimate the values at uniform k/N locations. At no point do we interpolate our k-space, making our algorithm distinct from regridding methods. Instead, we modified (14) to account for irregularly sampled k-space, while leaving the image space discrete, making it a special case of a nonuniform Fourier transform.

For our purposes, it is more convenient to express the k coordinates *k*_*x*_ and *k*_*y*_ not as discrete variables, but as an array of *k*_*x*_, *k*_*y*_ coordinates in the order in which they were sampled. We define these arrays as *k*_*x*_(*t*) and *k*_*y*_(*t*), which are given by equation (6). Essentially, *k*_*x*_(*t*) is the true *x* coordinate of k-space acquired at time t. Likewise, *k*_*y*_(*t*) is the true *y* coordinate of k-space acquired at time t. In this format, plotting *k*_*x*_(*t*) vs *k*_*y*_(*t*) would trace the k-space sampling trajectory, like those shown in Figure 2 (a), (c), and (e). We also add a phase compensation factor to account for translation induced phase shifts, shown in equation (15):

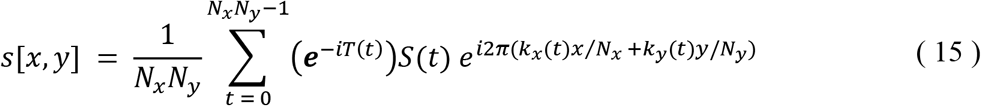

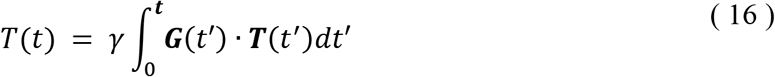

where and *S*(*t*) is raw fMRI data at coordinates *k*_*x*_(*t*) and *k*_*y*_(*t*). If there is no translation, *T*(*t*) equals zero, and so the exponential factor reduces to 1. The full equation reconstructs irregularly sampled k-space trajectories into discrete image space. The “discrete reconstruction of irregular fMRI trajectories” (DRIFT) algorithm can be applied directly to motion-contaminated data.

If the rotation R(t) is known at every dwell time, (6) can be used to calculate *k*_*x*_(*t*) and *k*_*y*_(*t*). One way to achieve this would be to use a high-speed motion capture camera, which could record the subject’s position at every point in time. This could then be used retrospectively to reconstruct the raw data when time is not an issue. However, the dwell time in a typical fMRI acquisition is anywhere between 5-12μs, which would require around 200k fps, which is well beyond the capabilities of MR compatible video cameras. If motion cannot be measured at every dwell time, we are still able to estimate the k-space coordinates at each dwell time if motion is sampled at a reasonably fast rate. In order to determine how fast the motion must be sampled for our estimations to be accurate, we examine the motion profile of over 6,000 scans, and found that 99% of frame-wise motion is between +/− 0.5mm for translation, and +/− 0.5 degrees for rotations. Even for the extreme outliers, the maximum rotation found was 7 degrees per volume. The primary concern with sampling such high rotations is the smoothness of the movement between samples. Typically, biological movements are relatively slow. One of the fastest motions a human can make is a blink of the eye, and even this lasts around 200ms. As head motion is slower than this, we determine that measuring the motion once per slice (usually around 60ms) is more than enough to accurately estimate the motion using linear interpolation. This corresponds to a frame rate of less than 20 fps, which is easily achievable for any MR compatible video camera. In fact, many cameras are capable of recording over 200 fps, which would even further improve our motion estimates.

Because of the relatively slow nature of biological movements, we can assume that any motion during slice acquisition is linear. For example, to model in plane rotation, if *k*_*x*_(*t*) and *k*_*y*_(*t*) are the intended, motion-free k-space coordinates, the motion contaminated coordinates can be estimated as:

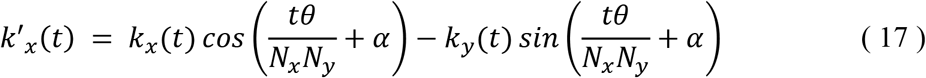

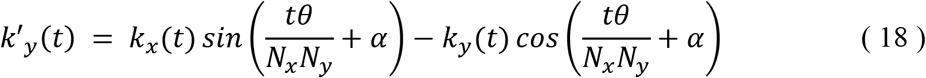

Where *α* represents any bulk rotation that has already occurred before k-space sampling (Accounting for offsets as in Figure 2 (e)). This evenly distributes a rotation of angle θ over N_x*_N_y_ time points, thus making *k*′_*x*_(*t*) and *k*′_*y*_(*t*) an incrementally rotated version of the intended trajectory, *k*_*x*_(*t*), *k*_*y*_(*t*). A similar procedure can be used to calculate the k-space coordinates for any rotation around the *x* and *y* axis.

By using either the estimated k-space coordinates from equations (17) and (18), or the true k-space coordinates from equation (6), we are able to completely recover irregularly-sampled k-space images for all plausible motions expected to be found in a typical fMRI scan. This algorithm is extremely versatile in its application. Not only can it be applied on a slice by slice basis to correct for k-space motion artifacts, but it can also be used to correct for bulk in-plane motion artifacts to realign slices and volumes without the need for interpolation. We now rigorously test the performance of our method in a variety of realistic fMRI conditions, using both real and simulated data. We use FSL’s Physics-Oriented Simulated Scanner for Understanding MRI (POSSUM) (Drobnjak *et al.*, 2006; Drobnjak, Pell and Jenkinson, 2010), a Bloch-equation based fMRI simulator to confirm the theory section of this paper. We examine the effects of various in-plane and out-of-plane motions on the fMRI signal, and the ability of DRIFT to remove them. Finally, we use a rotating phantom to test the application of our method on 3T Siemens Prisma scanner.

## Methods

### Simulated data

POSSUM requires a set of acquisition parameters, a full pulse sequence, and a target object with a defined geometry and spatially varying T_1_ recovery and T_2_ relaxation rates as inputs. The POSSUM simulator then takes the pulse sequence and target image as inputs, and generates an MR signal by simulating the proton spins in a static magnetic field, and solving the Bloch equation at every point in time during the scan. POSSUM also simulates rigid body motion of the object for any given motion parameters, which carries out motion at every update of the Bloch equation, allowing for the simulation of motion during readout and RF excitation.

Our simulated fMRI data in this study consisted of 21 brain volumes, each with 7 slices, with simulated motion occurring during the acquisition of the center slice (slice 4) in volume 10 for all cases. Our target object was POSSUM’s default high resolution brain image (voxel size 1mm × 1mm × 1mm, FOV 181mm × 217mm × 181mm), containing tissue segmentation for white matter, grey matter, and CSF. Each tissue type has a pre-defined T_1_ recovery, T_2_ relaxation, and proton density value: (Grey matter: T1 = 1.331s, T2 = 0. 051s, ρ = 0.87. White matter: T1 = 0.832s, T2 = 0.044s, ρ = 0.77. CSF: T1 = 3.7s, T2 = 0.5s, ρ = 1.0). For all simulations, the read direction was aligned with the *x* axis, and phase encoding along *y*. Slice selection was performed along the *z* axis. All parameters were simulated in a 3T magnetic field. Noise is added to the simulations to represent a realistic scan. POSSUM simulates system noise as independent, additive white Gaussian noise in the receiver channels, so the noise is present in the k-space before reconstruction. We chose a standard deviation of 0.01658 (units of intensity) to match the temporal SNR found in the real data acquired for this experiment, defined as the mean temporal signal divided by the standard deviation of the noise (commonly called tSNR) (Triantafyllou *et al.*, 2005). No B_0_ inhomogeneities or chemical shifts were added to the simulation to better focus on the effects of motion. The fMRI scan dimensions were 112×112×7 voxels, with a voxel size of 2×2×4mm, giving a 224×224×28 FOV. Our simulated pulse sequence had a TE of 60 ms, a TR of 1.0s, and a 60 degree flip angle. We choose a higher TE than most fMRI pulse sequences to magnify motion artifacts for the purpose of highlighting how motion effects the fMRI signal.

We chose 10 different kinds of motion with two different motion profiles to simulate in POSSUM. These motions are categorized as in-plane and out-of-plane and listed in table 1: 1) 5 degree rotation around *z*. 2) 5mm translation along *x*. 3) 5mm translation along *y*. 4) 5mm translation along both *x* and *y* together. 5) 5mm translation along *y* with a 5 degree rotation around *z*. 6) 5 degree rotation around *x*. 7) 5 degree rotation around *y*. 8) 5mm translation along *z*. 9) 5 degree rotation around *y* with PMC, and 10) 5 mm translation along *z* with PMC. Scans 1-5 simulate “in-plane” motion, meaning the motion occurs in a way so that the excited tissue remains within the plane of acquisition, and does not cause any disruption to the steady-state. These scans demonstrate DRIFT’s ability to remove k-space motion artifacts from in-plane motion. Scans 6-8 demonstrate out-of-plane motion with the harmful effect of spin-history artifacts, and scans 9-10 simulate PMC and demonstrate that if spin-history artifacts are avoided, DRIFT can again correct for any readout motion artifacts that are present.

**Table 1:** 10 motion profiles used for simulated data.

| Simulation Type | Sim # | Motion Type | Motion Profile | Rx | Ry | Rz | Tx | Ty | Tz |
| --- | --- | --- | --- | --- | --- | --- | --- | --- | --- |
| In-plane | 1 | Z-axis Rotation | *a* | - | - | 5° | - | - | - |
|  | 2 | X-axis Translation | *a* | - | - | - | 5mm | - | - |
|  | 3 | Y-axis Translation | *a* | - | - | - | - | 5mm | - |
|  | 4 | X and Y-axis Translation | *a* | - | - | - | 5mm | 5mm | - |
|  | 5 | Z-axis Rotation and Y-axis Translation | *a* | - | - | 5° | - | 5mm | - |
| Out-of-plane | 6 | X-axis Rotation | *a* | 5° | - | - | - | - | - |
|  | 7 | Y-axis Rotation | *a* | - | 5° | - | - | - | - |
|  | 8 | Z-axis Translation | *a* | - | - | - | - | - | 5mm |
|  | 9 | Y-axis Rotation | *b* | - | 5° | - | - | - | - |
|  | 10 | Z-axis Translation | *b* | - | - | - | - | - | 5mm |

The profiles of the simulated motion relative to the pulse sequence are shown in Figure 3. The RF pulse, *z*, *y*, and *x* gradients are depicted in the first 4 lines, respectively. The 5^th^ line, labeled “Mot”, refers to the relative displacement of the object, and can in general refer to any type of motion. The line can indicate a translation, rotation, or combination of the two along any or multiple axis, and is intended to demonstrate the relative timing of the motion in the pulse sequence. In Figure 3 (a), motion begins at the first point of k-space sampling, and continues linearly until the end of slice acquisition. The target then remains at its final position for the remainder of the scan. This is referred to as motion profile “a”, and would result in misalignment, as well as spin-history artifacts if out-of-plane motion is present. The second motion profile, shown in Figure 3 (b), begins the same as Figure 3 (a), where motion starts at the beginning of k-space sampling and ends and the end of readout. However, in this profile, the object returns to its original position before the next RF pulse, as indicated by the “Mot” line returning to zero. This would not cause spin-history artifacts, regardless of the type of motion. This is referred to motion profile “b”, and is equivalent to adjusting the gradients to account for subject motion, as in PMC.

**Figure 3:**
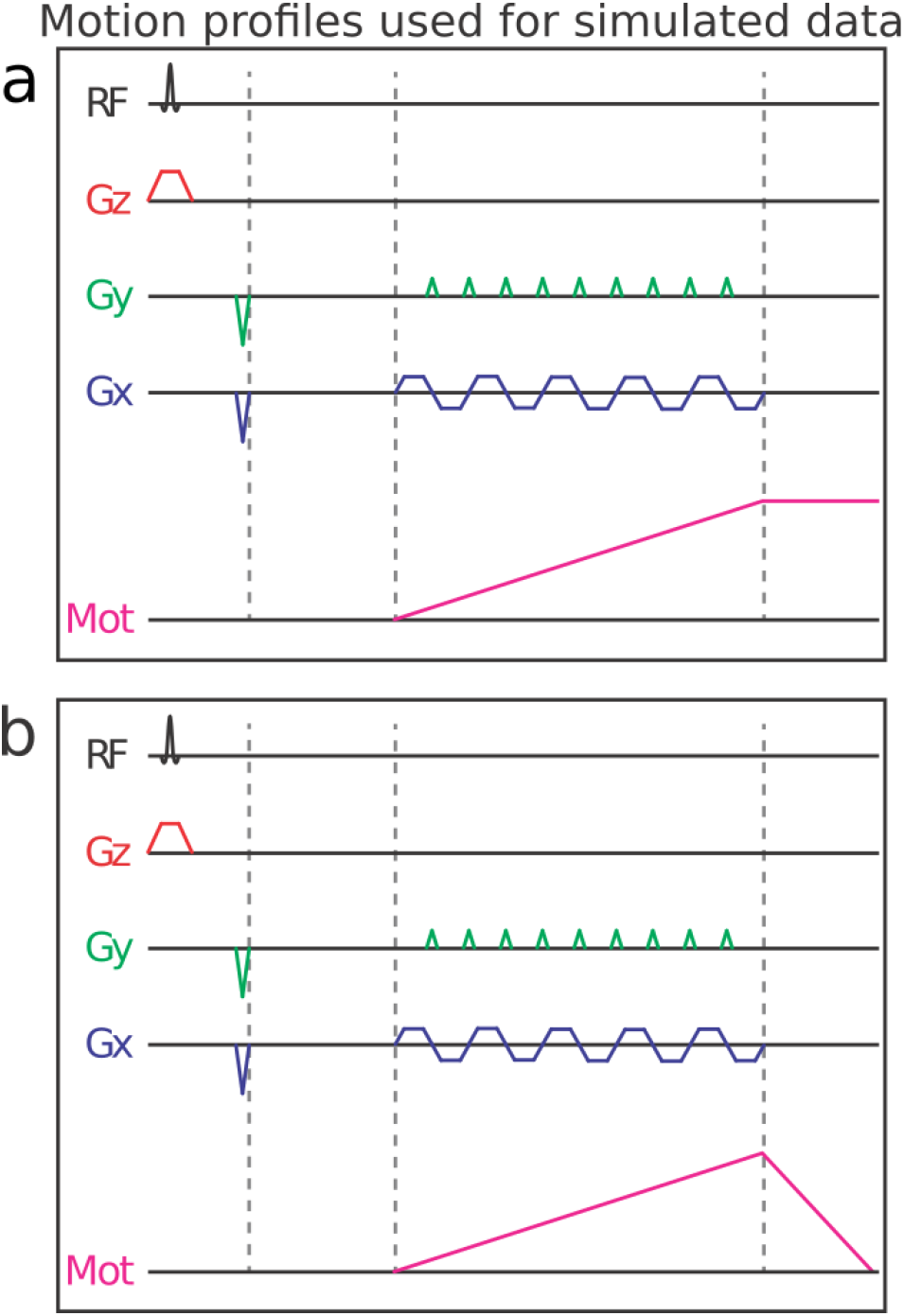
An illustration of the motion profiles used in simulated data. (a) The motion begins after RF excitation at the start of k-space readout. Motion ramps continuously until the end of k-space readout, at which point the object stops moving and remains at its new position. (b) The motion begins as in (a), after RF excitation and at the start of k-space readout. Motion ramps continuously until the end of k-space readout, at which point the object quickly returns to its original position before the next RF pulse.

While each of the above simulations were created with exaggerated movements to examine the effects of different types of motion on an fMRI image, it is important to also examine the effect of real subject motion in our simulation. Therefore, another simulation consisting of 41 volumes is performed with similar acquisition and target parameters as our previous simulations, but with in-plane rotation and translation parameters estimated from a real subject motion during a task-based fMRI scan (mean framewise rotation = 0.4 degrees, and mean framewise translation = 0.3mm). 36 in-plane rotation measurements were used from this simulation, and the first 5 volumes were simulated with no motion to reach the steady-state, and volume 5 was used as a reference volume. We use this simulation to demonstrate the effectiveness of DRIFT on a scan with more realistic motion, rather than the large, isolated movements used in our previous simulations. Each rotation/translation occurs over the length of a full TR, rather than a single TE, as in the previous simulations.

### Real Data

An in-house MRI-compatible phantom capable of in-plane rotation was developed and used to acquire our real data using a Siemens 3T Prisma scanner. The phantom consists of two parts: an imaging portion which has identifiable geometry and can be filled with agar gel, and a driver portion, which includes fan blades and can be rotated with air pressure in a controlled manner. We designed the phantom using FreeCAD, and fabricated the structure using a Formlabs Form 2 stereolithography 3D printer. The phantom is a hollow cylinder 2.25 inches tall with a 4 inch diameter. The phantom was printed with photoreactive resin RS-F2-GPCL-04 consisting of a proprietary mixture of methacrylic acid esters and photoinitlators. Several geometric shapes protrude from the base of the phantom to provide contrast for scanning, shown in Figure 4. An agar solution was created and poured into the phantom and allowed to solidify. The agar solution was 15% salt by weight to prevent bacterial growth on the agar gel. The phantom was mounted on the driver portion to allow for rotation. A pressure regulator was used to maintain a relatively constant PSI which allowed a controlled rate of rotation. Rotation rate was measured through manual realignment of EPI scans. A full depiction of our setup is shown in Figure 5.

**Figure 4:**
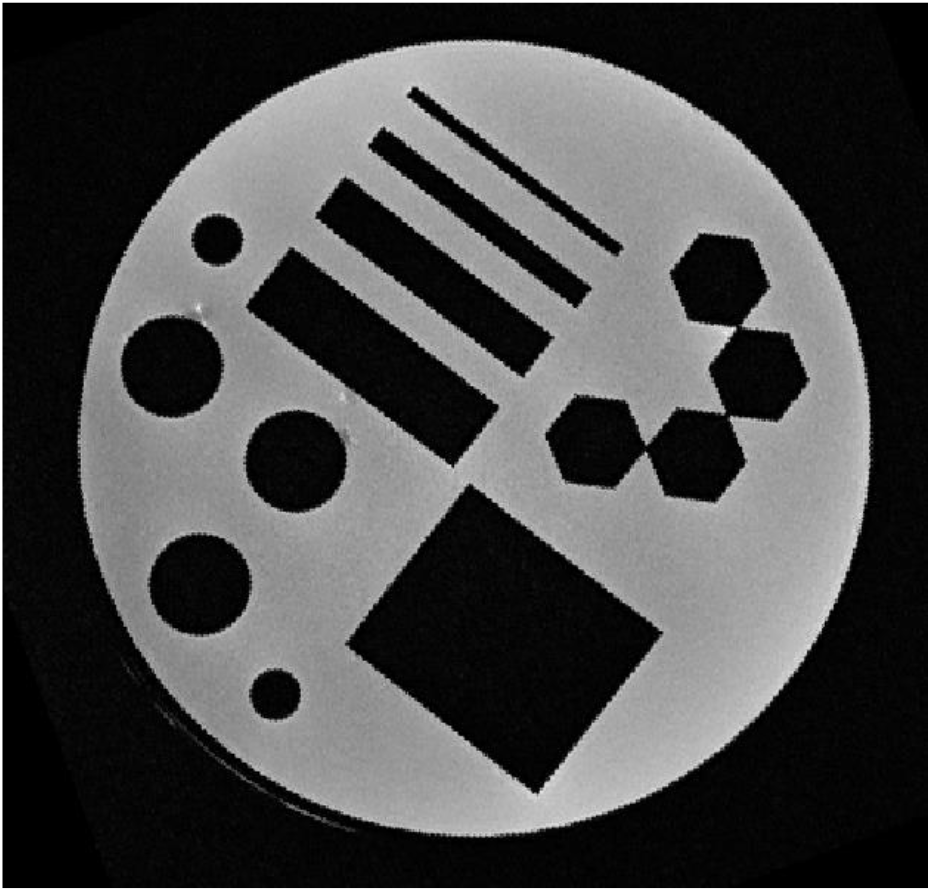
A high-resolution T2 image of the agar filled phantom used for real data scans.

**Figure 5:**
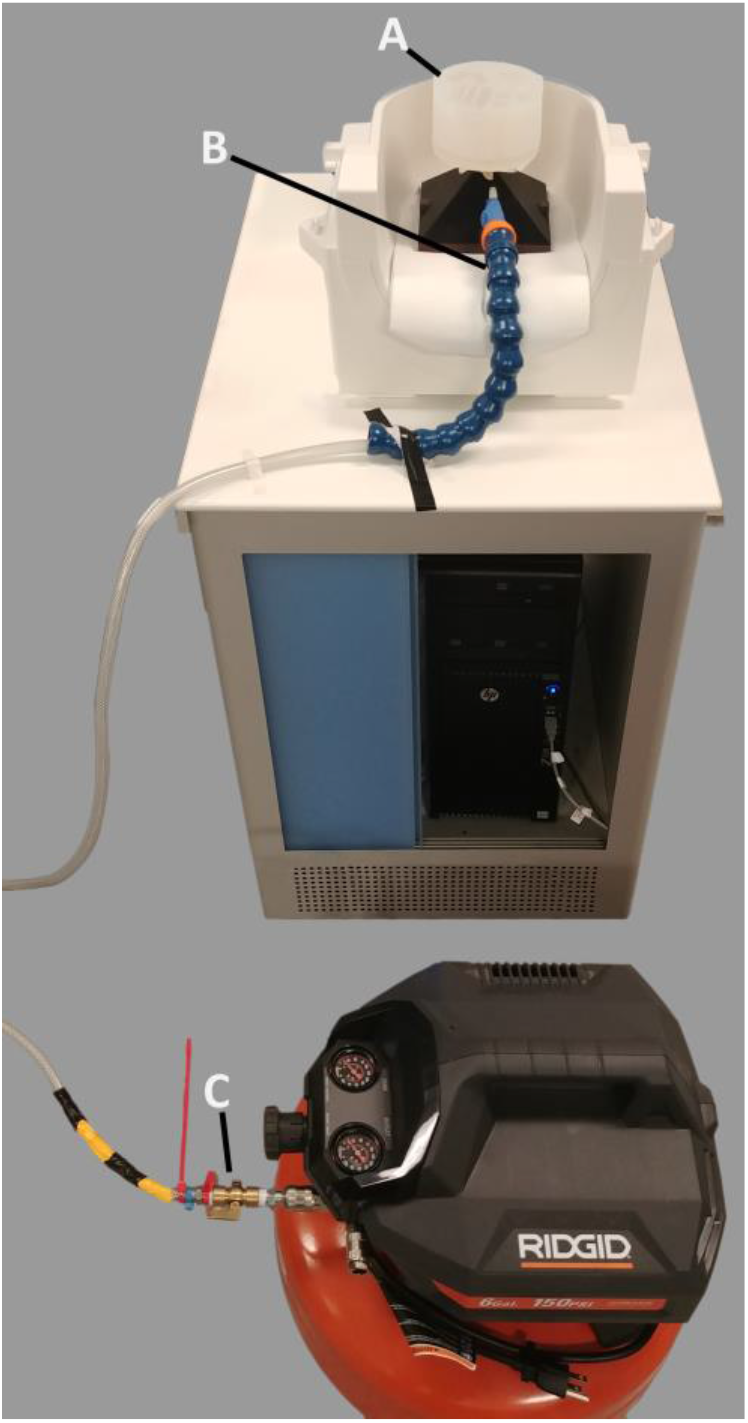
Setup of the rotating phantom. (A) The imaging section of the phantom rests atop its rotating stand in a head coil. (B) A directional nozzle focus air flow onto the fan blades in the rotating stand. (C) The pressure regulator and flow control valve which allow us moderate control over the degree of rotation.

We began the scan with 10 seconds of motionless volumes to use as a reference and to calculate SNR at baseline. Then, the airflow was turned on to allow moderate rotation to take place over the next 40 seconds. The airflow was shut off in time to allow the phantom to come to rest for the final volumes of the scan. The framewise rotations for our scan ranged between 1 and 130 degrees with a median rotation of 54.5 degrees (0.15 hz). One coronal slice was acquired with a 64×64 matrix, with an FOV of 127×127×2mm. We used a basic EPI pulse sequence, with a TR/TE of 1.0s/30ms, flip angle 90, with no slice acceleration. The phantom was placed inside a 64 channel head coil which was used for image acquisition. We acquired 60 volumes, the first 10 of which were stationary, followed by 38 volumes of motion, and an additional 12 volumes at the end of the scan with no motion, but at a random angle offset from the initial orientation. Both the Siemen’s reconstructed DICOM images, as well as the raw data were acquired and used in processing.

### Simulated Data Processing Pipeline

The reconstruction provided by POSSUM is contaminated with k-space motion artifacts and misalignment, and is referred to as the “uncorrected” data, meaning no motion correction has been applied. We then use fMRI realignment software to perform rigid body realignment on the data. This is referred to as the “realigned” data set. Because the motion parameters are known, we are able to provide precise rotations and translations to the realignment software, eliminating any possible artifacts due to inaccurate motion estimation. For the volume in which motion occurs in, half of the slices are acquired normally, then motion is applied during one slice, and the remaining half of the slices are acquired with misalignment. This is problematic for rigid body registration, as half the brain is misaligned. For this volume, the average motion across all slices was used to realign the volume. Finally, we create the “DRIFT” reconstruction using the provided pulse sequence, known motion parameters, and raw data. We calculate the effect of motion on the k-space trajectory using custom python code that evaluates equations (6) and (7). We then use these true k-space coordinates to reconstruct the raw data using DRIFT.

While most modern MRI scanners discard initial volumes that are not in steady-state, POSSUM leaves the scans in the simulation. These volumes have a different intensity and contrast than the later volumes that have reached steady-state. We discard the initial 5 volumes from the simulations, at which point the voxel’s have settled to 99.99% of their steady-state value. After reaching the steady-state, we do not expect to detect any change in the reconstructed images except the additive white Gaussian noise. Any further change in the signal should be attributed to the motion. We quantify the effect of motion for each reconstruction by computing the percent difference (PD) between each motion contaminated/corrected slice and the uncontaminated slice acquired in volume 8, henceforth referred to as the reference volume. Percent difference is computed by following equation.

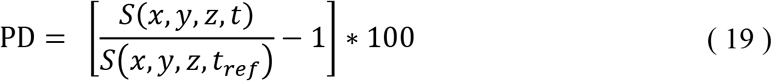

Where t_ref_ is the timepoint of the reference volume to be used as a baseline. We then calculated the PD for a motionless volume (“Before Motion”, volume 9), the volume in which the motion occurred (“During Motion”, volume 10), and for the volume acquired immediately after motion (“After Motion”, volume 11). Any error in the “Before Motion” volume is due to noise, while any additional error in the “During Motion” volume is due to motion artifacts during slice acquisition. Finally, any changes in the volume after motion are due to bulk motion and/or spin-history artifacts. For the 10 scans examining exaggerated subject motion, PD is calculated for slice 4, which is the slice that motion occurs in, and will have the artifacts associated with k-space displacement. For the simulation with real subject motion, we chose volume 5 to be used as a reference volume (as all previous volumes are not in steady-state, and all following volumes have motion), and calculate the PD over the entire volume, since motion now occurs over the entire volume rather than one single slice. 95% confidence intervals are bootstrapped for the PD’s of each reconstruction method and shown in the resulting analysis (Dragicevic, 2016). Statistical inference is done on voxelwise PD values using a repeated measures two-sided t-test.

### Real Data Processing Pipeline

There are very few tools available to process raw k-space data, and given that our algorithm must be implemented in place of a normal IFFT reconstruction, it was necessary to create our own python code to reconstruct and process the images. First, the raw k-space data from all 64 channels is read into the processing pipeline where noisy channels are removed by examining the correlation of each channel with all other channels. A correlation below 0.5 was used as an exclusion threshold, which occurs only when the SNR is extremely low. Upon inspection, the channels omitted with this process had little or no recognizable image in its reconstruction. We then apply a small hamming window filter on the high-frequency k-space components to suppress high frequency noise, which is a common step in many k-space reconstruction routines (Lowe and Sorenson, 1997; Friedman *et al.*, 2006; Caparelli and Tomasi, 2008). The window is applied to the 14 outermost k space samples in the *x* and 7 outermost k-space samples in the *y* direction, as Siemen’s oversamples in the read direction (*x*) by a factor of 2. We then use the calculated k-space coordinates with the DRIFT algorithm to reconstruct two images, one image from the odd read samples and one from the even read samples. The average of these two images is stored as the reconstructed image for a single channel. The channels are then combined using a root sum of squares method. We also create a second reconstruction, using identical processing steps with the exception of the motion correction portion of DRIFT. This data by itself is not corrected for motion in any way, and is referred to as the “uncorrected” data set. We then run realignment on this data, using the same method described for simulated data, to evaluate the effectiveness of traditional realignment methods. This leaves us with three reconstructed scans: Uncorrected, realigned, and DRIFT corrected data. This gives us a direct comparison on the effect of DRIFT on the reconstructed data, with no additional confounds in the preprocessing stream.

To quantify the amount of motion at each volume, we used the scanner reconstructed images to manually estimate motion between volumes. The first volume was chosen as a reference. The absolute rotation of each volume relative to the reference was estimated visually using Freeview. We also measured the relative rotation of each volume to the previous volume. The relative rotation to the previous volume gives the total rotation over one TR. Dividing this rotation by the TR gives an estimate of the rate of rotation in degrees per second for that volume. This rotation is used to estimate the true locations of k-space samples as in equations (17) and (18). In the EPI pulse sequence implemented by Siemens, signal acquisition occurs during the ramp up, plateau, and ramp down portion of the readout gradient. This leads to non-uniformly spaced k-space samples. Siemen’s EPI reconstruction will use some form of regridding to compensate for this ramped sample trajectory, however since our method is already able to account for irregular sampling offsets, we simply use the true, nonuniform k-space points, and apply the appropriate rotation directly to them. An example of the sampling trajectory for a single read line is stored in the raw data header, which we extract and use to generate our k-space.

Due to intensity distortions and non-uniformity in the rotating reconstructed images, we could not use PD as our measurement for quantitative evaluation of DRIFT. Instead we calculate the mutual information between each volume in each reconstruction and a baseline volume. We generate our baseline volume by averaging the first 10 motionless scans. DRIFT is able to reconstruct the raw data without introducing any interpolation error or smoothing to the reconstructed image, whereas interpolation of any kind contains an inherent low-pass filter effect, causing a slight smoothing of the image. It is well known that smoothing can inflate or induce correlations between two images or signals (Friston *et al.*, 1995; Davey *et al.*, 2013; Razlighi *et al.*, 2013). For a fair comparison, we calculate the effective equivalent Gaussian smoothing kernel to replicate the smoothing induced by realignment. To do this, we extracted the final volume from the realigned scan, and calculated its correlation with the first volume in the uncorrected scan. The uncorrected scan was then smoothed with Gaussian kernels of increasing size, incrementing the FWHM by 0.1mm. We plot the correlation as a function of the smoothing kernel’s FWHM in Figure 6 (a). The maximum correlation is taken as the effective FWHM of interpolation, corresponding to 2.38mm, or a standard deviation of 1.01mm. Upon visual inspection, the smoothed image was visually indistinguishable from the FLIRT corrected image, as shown in Figure 6 (b) and (c). We applied this kernel to both the uncorrected and the DRIFT corrected data to remove smoothing-induced correlation as a confound. Similar to the simulated data, statistical inference was carried out as a repeated measures two-sided t-test on mutual information measurements.

**Figure 6:**
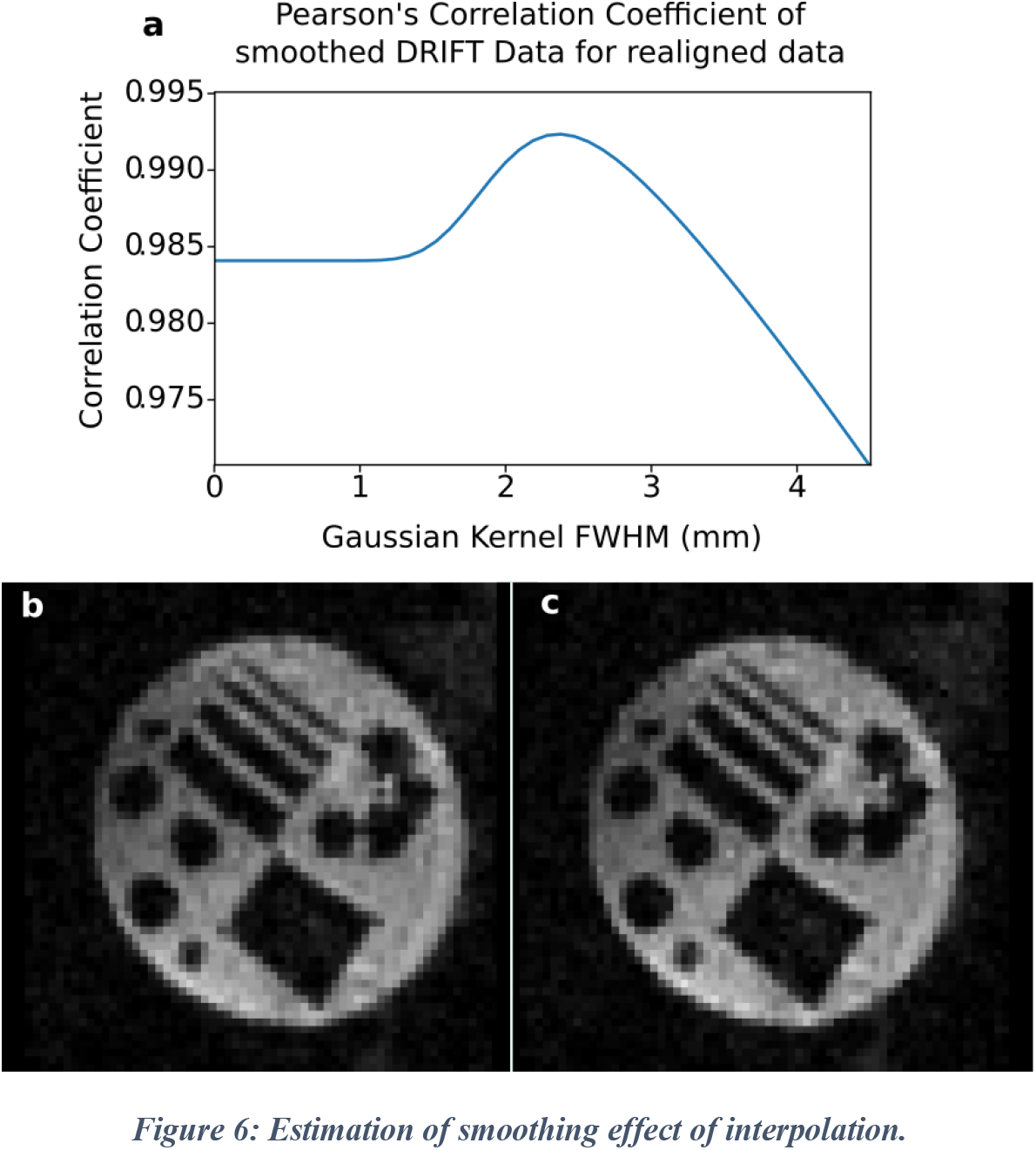
Estimation of smoothing effect of interpolation. (a) A stationary but misaligned volume is selected, and it is realigned to a reference slice. The reference slice is then smoothed with progressively larger Gaussian kernels, and the correlation is calculated between the realigned volume and the smoothed reference at each iteration. A maximum correlation was found with a Gaussian FWHM of 2.3mm. (b) Volume 48 of the DRIFT corrected data with a 2.3mm FWHM smoothing kernel. (c) Volume 48 of the realigned volume, with no smoothing other than that inherent in interpolation.

## Results

We first use simulated fMRI data to demonstrate that each type of motion generates strikingly different types of artifacts. Next we use data collected from a rotating phantom to establish the proof of concept for our proposed method of hybrid motion correction.

### Effects of in-plane motion and its correction

In-plane motion does not disturb the steady-state of the EPI sequence, thus they will not cause any spin-history artifacts. Figure 7 illustrates the results of our simulation for in-plane motion. In this simulation, during the acquisition of one slice (slice number 4) in volume 10, a 5 degree rotation around *z* and/or 5mm translation along the *x* or *y* axes is applied to the target object. The top row in Figure 7 depicts the artifact of each type of in-plane motion on the reconstructed image using simulated data with an exaggerated motion profile. To make the artifacts visually identifiable, the simulated motion in this row is three times larger than the motion used in bottom 2 rows. These exaggerated simulations are used only in the top row for visualization, and all statistical comparisons are carried out on the data described in the methods section. The blurring on the boarder of the brain and ringing effects inside the brain are visible in this row. Furthermore, a severe geometric distortion is also visible particularly for the combination of translation and rotation. To investigate the actual differences in the fMRI signal, we computed the PD, as defined in equation (19).

**Figure 7:**
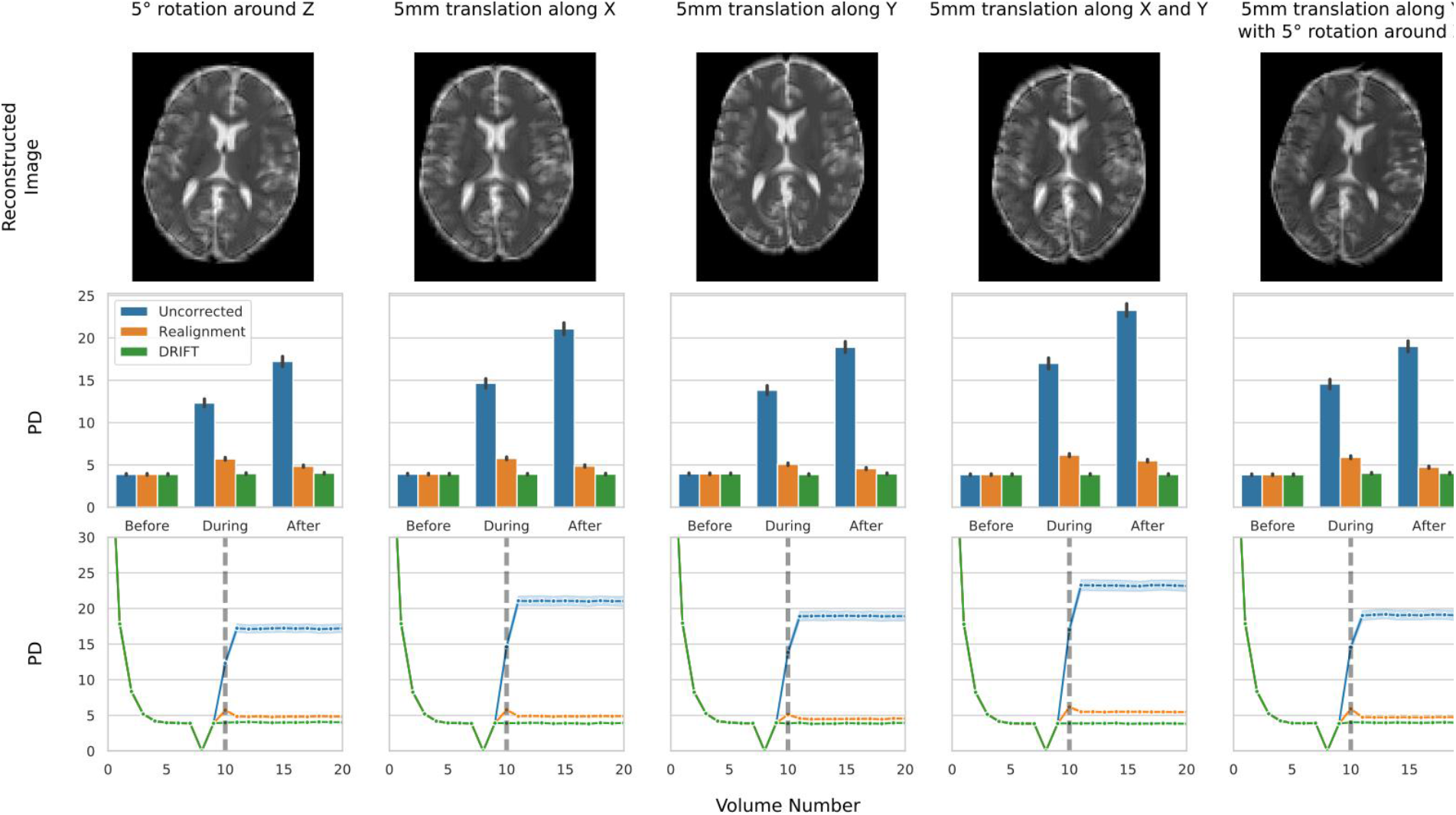
Artifacts and PD for in-plane motion, in simulated data. (Top) Illustration of the artifacts that occur for different types of in-plane motion. (Middle) PD calculated for the slice in which motion occurs for the volume before, during, and after motion. (Bottom) Timeseries of the PD calculated for the slice in which motion occurs for the entire simulation. Volumes 9, 10, and 11 correspond to the before, during, and after volumes respectively, used in the middle row.

To investigate the actual differences in the fMRI signal, we computed the percent difference (PD), as defined in method section, between the selected slice (slice 4) in a reference volume (volume 8) and the slice acquired immediately before applying motion (in volume 9), the slice acquired during the motion (in volume 10), and the slice acquired immediately after motion (in volume 11). The middle row in Figure 7 shows the PDs for uncorrected, realigned and DRIFT corrected data. Before motion, there is a constant 3.8 PD across all three methods and for all five types of in-plane motion. This is due to the additive white Gaussian noise in our simulated data, and represents the lowest possible error in a motionless volume. Motion will introduce error and artifacts into the reconstruction, which will raise the PD above this value. Successfully correcting for motion will bring the PD back down to this baseline value.

The difference between uncorrected, realigned and DRIFT corrected slices became significant for the slice acquired during motion. As expected, realignment significantly reduced the motion artifact depicted in top row in comparison to the uncorrected slices (PD_diff_=6.6, p < 10^−5^, for 5 degrees rotation around z; PD_diff_=8.9, p < 10^−5^, for 5mm translation along x axis; PD_diff_=8.7, p < 10^−5^, for 5mm translation along y; PD_diff_=10.8, p < 10^−5^, for 5mm translation along x and y; PD_diff_=8.7, p < 10^−5^, for 5 degrees rotation around z and 5mm translation along y). However, there are still remaining artifacts in all five cases of in-plane motions. Using DRIFT not only significantly reduces the PD between the realigned and DRIFT corrected slices (PD_diff_=1.7, p < 10^−5^, for 5 degrees rotation around z; PD_diff_=1.9, p < 10^−5^, for 5mm translation along x; PD_diff_=1.2, p < 10^−5^, for 5mm translation along y; PD_diff_=2.3, p < 10^−5^, for 5mm translation along x and y; PD_diff_=1.9, p < 10^−5^, for 5 degrees rotation around z axis and 5mm translation along y), but also the improvement reached the point that there was no significant difference between the motionless slice and DRIFT corrected slice during motion (p > 0.05 for all motions), suggesting that DRIFT can potentially remove the motion artifact completely from the motion contaminated slices acquired during any in-plane movement. After motion, the PD for the uncorrected slice increases compared to the slice acquired during motion. This can be due to the fact that motion contamination accumulates gradually over the k-space filling; as shown in Figure 2 (c), there is a negligible motion artifact in the beginning of k-space filling, whereas it becomes most dominant at the end. However, the slice acquired in the next volume is completely rotated/translated causing an increase in the PD for the uncorrected slice. While realignment should correct for post motion displacement, the interpolation artifact during the spatial transformation prevents the PD from reaching its baseline value before motion, causing a significant difference between the realigned and DRIFT corrected slice (PD_diff_=0.8, p < 10^−5^, for 5 degrees rotation around z; PD_diff_=1.0, p < 10^−5^, for 5mm translation along x; PD_diff_=0.6, p < 10^−5^, for 5mm translation along y; PD_diff_=1.6, p < 10^−5^, for 5mm translation along x and y; PD_diff_=0.7, p < 10^−5^, for 5 degrees rotation around z and 5mm translation along y). This results highlight the superiority of the DRIFT motion correction technique even for simple instantaneous transformation compared to conventional spatial transformations.

The bottom row in Figure 7 shows the computed PD for slice number 4 in all 20 simulated volumes in uncorrected, realigned, and DRIFT corrected data. The dots on the curve mark the mean of the PD and the shaded band around each curve illustrate the 95% confidence interval at each point of computation. As demonstrated by this figure, all three scans reach their steady-state by volume five, and drop to zero at volume 8 (since that is the selected reference for computing PD). They return to their steady-state value at volume 9. The motion contaminated slice in volume 10 demonstrates an abrupt increase in PD in both uncorrected and realigned slices whereas the DRIFT corrected slice remains unchanged. In volume 11, after the occurrence of motion, the uncorrected slice reaches its highest PD and plateaus from that point on, whereas realignment slightly but significantly reduces the PD of the slice acquired after motion and remains constant afterward. DRIFT corrected slices have unchanged PD that remain identical to the PD in “before motion” volumes, indicating a complete removal of in-plane motion artifacts.

To summarize, uncorrected data has the largest PD for all during and after motion volumes. Realignment significantly reduces this error, but is unable to account for k-space filling artifacts, and still has significant interpolation error in the volumes after motion. DRIFT significantly reduces the PD beyond what realignment can do in all cases, and all DRIFT reconstructed volumes are not significantly different from the motionless baseline.

### Effects of out-of-plane motion and its correction

As discussed in the method section, out-of-plane motion disturbs the steady-state of the EPI acquisition, thus causing spin-history artifacts which has a lasting effect up to a few volumes after the occurrence of motion. Figure 8 illustrates the results of our simulation for out-of-plane motion. In this simulation, during the acquisition of one slice (slice number 4) in volume 10, we show the effect of a 5 degree out-of-plane rotation around *x*, a 5 degree out-of-plane rotation around *y*, as well as a 5 mm translation along *z*. In addition, in order to simulate the effect of prospective correction applied simultaneously with DRIFT, we rerun the simulations of a 5 degree rotation around *y* axis and a 5 mm translation along the *z* axis using the motion profile shown in Figure 3 (b), and applied a quicker inverse motion to the target object to come back to its original location before application of next slice selection RF pulse.

**Figure 8:**
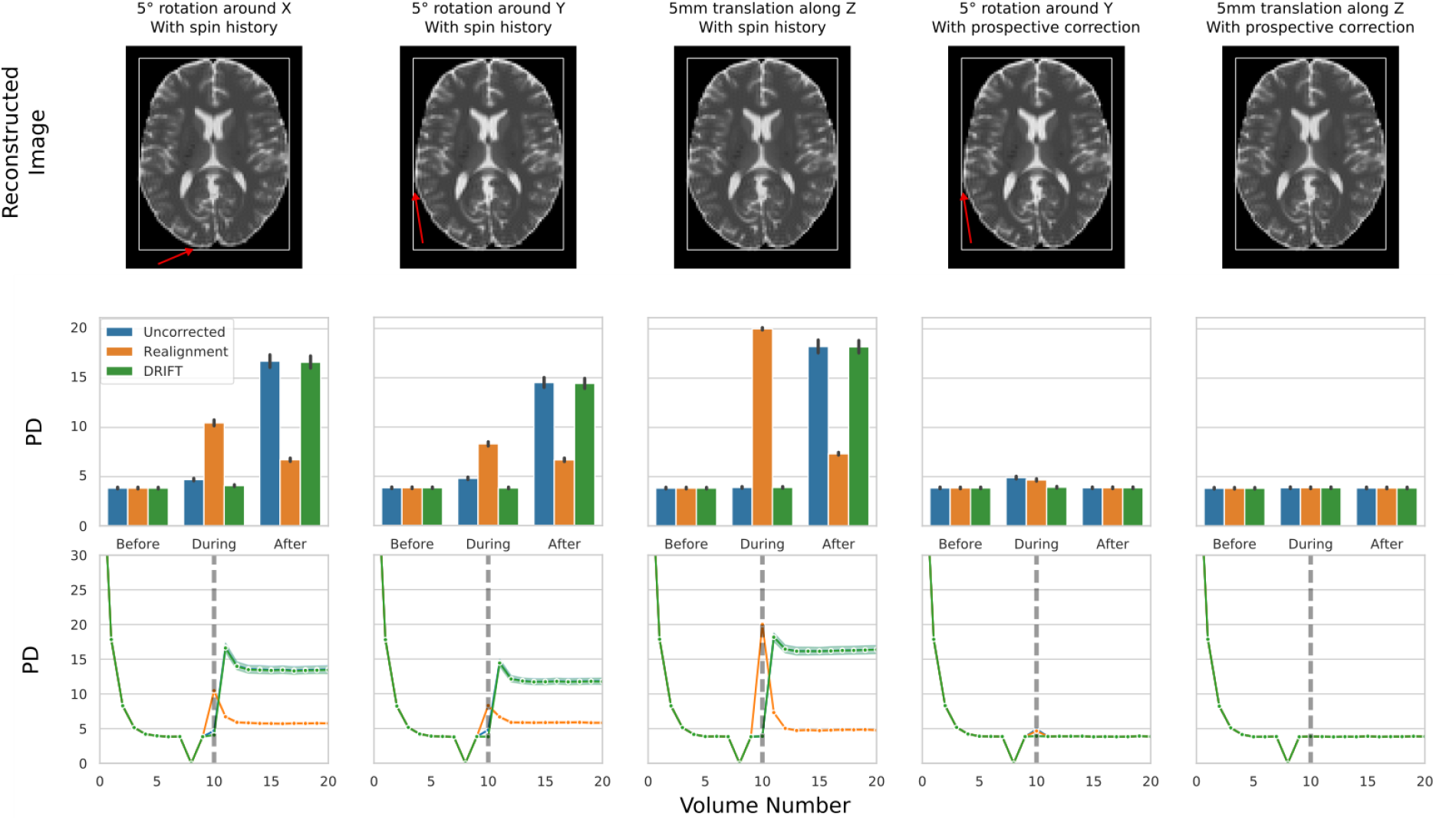
Artifacts and PD for out-of-plane motion, in simulated data. (top) Illustration of the artifacts that occur for different types of out-of-plane motion. (middle) PD calculated for the slice in which motion occurs for the volume before, during, and after motion. (bottom) timeseries of the PD calculated for the slice in which motion occurs for the entire simulation. Volumes 9, 10, and 11 correspond to the before, during, and after volumes in the middle row.

The top row in Figure 8 depicts the artifact of each type of motion on the reconstructed image using simulated data with an exaggerated motion profile. To make the artifacts visually identifiable, the simulated motion in this row is three times larger for rotation and translations. These exaggerated simulations are used only in the top row for visualization. A white box is placed around each brain volume that is the exact dimensions of a motionless slice. The artifacts due to out-of-plane motion are extremely small, but can be seen as a slight compression of the image, indicated by a small gap between the brain and the white box, highlighted by red arrows. Note that these artifacts are extremely small when present, and that there are no gaps for the *z* translation volumes. We have already demonstrated these results through equations (7), (11), and (12). In fact, according to our equations, the out-of-plane translation should not cause any artifact on the slice where the translation took place.

The middle row in Figure 8 shows the PDs for uncorrected, realigned and DRIFT corrected slice before, during, and after out-of-plane motion. Similar to in-plane simulations, before motion there is a constant 3.8 PD across all three methods and for all five types of out-of-plane motions, which is due to the additive white Gaussian noise. The difference between uncorrected, realigned and DRIFT corrected slices became significant for the slice acquired during motion. Out-of-plane rotations caused a slight but significant increase in the PD of the uncorrected slice in comparison to the motionless baseline (PD_diff_=0.9, p < 10^−5^, for 5 degrees rotation around x; PD_diff_=1.0, p < 10^−5^, for 5 degrees rotation around y). Interestingly, realignment makes the artifact even worse than the uncorrected slice (PD_diff_=5.8, p < 10^−5^, for 5 degrees rotation around x; PD_diff_=3.5 p < 10^−5^, for 5 degrees rotation around y). However, DRIFT completely removes the rotational artifact to a level that there is no significant difference between DRIFT corrected and the motionless baseline slices (PD_diff_=0.12, p > 0.05, for 5 degrees rotation around x; PD_diff_=0.0, p > 0.5, for 5 degrees rotation around y). For out-of-plane translation, as we predicted there is no significant difference between the translated and steady-state slice in uncorrected data (PD_diff_=0.1, p > 0.1). However, realignment again fails to correct for out-of-plane translation (PD_diff_=16.2, p < 10^−5^ compared to the motionless baseline), where the DRIFT corrected slice has no motion artifact (PD_diff_=0.1, p > 0.1 compared to the motionless baseline). This is mainly because realignment is a registration-based method that fails drastically for partially contaminated volumes.

As we discussed before, the dominant effect of out-of-plane motion takes place after the motion due to spin history. As shown in Figure 8 middle row, for uncorrected data the slice acquired immediately after the rotational and translational motions has a much higher PD compared to the slice where the motion actually took place (PD_diff_=12.0, p < 10^−5^, for 5 degrees rotation around x; PD_diff_=9.7, p < 10^−5^, for 5 degrees rotation around y; PD_diff_=14.3, p < 10^−5^, for 5 mm translation along z). While this is mostly due to misalignment, the PD of the realigned volumes are still significantly higher than the motionless baseline, despite being perfectly realigned (PD_diff_=2.9, p < 10^−5^, for 5 degrees rotation around x; PD_diff_=2.8, p < 10^−5^, for 5 degrees rotation around y; PD_diff_=3.5, p < 10^−5^, for 5 mm translation along z). This is because the spin history artifact contaminates the slice without causing any misalignment, thus making it impossible for a registration-based technique to correct for it. DRIFT also fails to correct for out-of-plane motion once that spin history artifact contaminates the slice, as it seen in the figure, the PD of the DRIFT corrected slice acquired after motion is significantly higher than the one acquired during motion (PD_diff_=12.5, p < 10^−5^, for 5 degrees rotation around x; PD_diff_=10.6, p < 10^−5^, for 5 degrees rotation around y axis; PD_diff_=14.3, p < 10^−5^, for 5 mm translation along z), suggesting that correcting for spin-history artifact is extremely challenging if not impossible.

While correcting for spin-history artifact seems to be a challenge, preventing it with slice-based prospective motion correction methods has already been proposed and implemented in the field. In the next two simulations, we aim to show that the combination of prospective motion correction and DRIFT, as a retrospective method, can optimally remove the motion artifacts from the fMRI data. By changing the motion profile and applying a quick inverse motion immediately after acquisition of the slice (and before applying next RF pulse), we aim to simulate our proposed hybrid motion correction technique. As shown in Figure 8, a 5 degree out of plane rotation around y again has a slight but significant effect on the PD of the slice where the motion took place in comparison to the motionless baseline (PD_diff_=1.0, p < 10^−5^, for uncorrected; PD_diff_=0.8, p < 10^−5^, for realignment), which is optimally corrected with DRIFT (PD_diff_ < 0.1, p > 0.1). For a 5 mm out-of-plane translation, no significant change has been observed in the fMRI data, if spin-history artifact has been prevented (PD_diff_ < 0.1, p > 0.5), essentially validating out findings in equation (7).

The bottom row in Figure 8 plots the computed PD for slice number 4 in all 20 simulated volumes with out-of-plane motion. Again, all simulated cases and correction methods reach their steady-state by volume 5. The dots on the curve show the mean of the PD and the shaded band around each curve illustrate the 95% confidence interval at each point of computation. The first three graphs represent motion with spin-history effects. For rotation around *x*, rotation around *y*, and translation along *z*, there is an abrupt increase in PD for the realigned slice at volume 10 (the volume in which motion occurs). However, we see only a small increase in PD for uncorrected data, and no increase in PD for DRIFT at volume 10. In volume 11, we now see a large increase in PD for both DRIFT and uncorrected data. At this point, we also see that the PD for realigned data has decreased slightly from volume 10. Over the next few volumes, the PD in all three data sets will gradually decrease until leveling out at a new baseline around volume 13. This gradual decrease is due to spin-history, and represents the system slowly returning to steady-state. The final two graphs repeat two of these motions, (5 degree rotation around *y* and 5mm translation along *z*) while preventing spin-history artifacts. In these simulations, there is no difference in the PD before and after motion for all correction methods, indicating a perfect removal of motion artifacts.

### DRIFT performance using real-subject motion parameters

Up to this point, all simulations were based on synthesized motion profiles which were exaggerated and performed mainly for the purpose of studying and differentiating the distinct effect each type of motion has on the data. In this experiment, we extracted motion parameters from a real subject’s typical fMRI scanning session and used it to simulate the effect of realistic motion on fMRI data. We used volume 5 as the reference volume where the fMRI signal reaches its steady-state equilibrium and computed volume-based PD to investigate the effect of realistic in-plane motion and evaluate the performance of our proposed motion correction technique. Figure 9 shows the results of this experiment. The three dotted black curves depict the in-plane motion parameters (rotation around *z* axis, transformation along *x* and *y* axes), and the solid black line indicates the displacement due to the motion parameters. The blue solid line indicates the PD computed for uncorrected fMRI volumes, and the orange and green curves shown the PD for realigned and DRIFT corrected volumes. The insert illustrates a magnified sections of the plots where there is minimal movement in the target object. The shaded border with the same color indicates the 95% confidence interval at each volume. Using realignment significantly attenuated the in-plane motion artifacts which is demonstrated by their PD curves.

**Figure 9:**
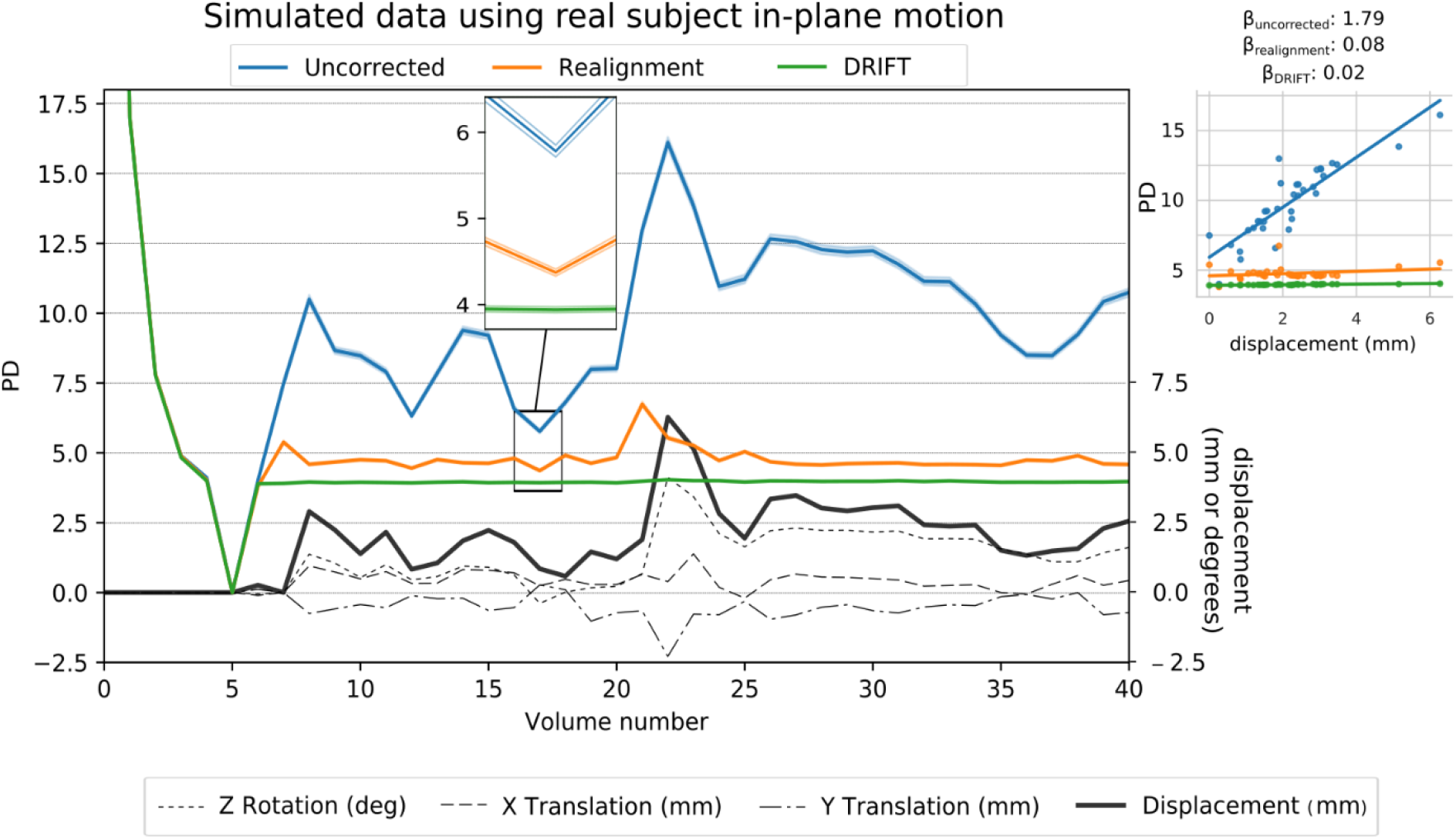
DRIFT correction on simulated data with real subject motion. Volume-wise absolute percent difference from baseline of 41 volume simulation with real subject in-plane rotation and translation for uncorrected, realigned, and DRIFT corrected data. Black dashed lines indicate the level of rotation, translation, and framewise displacement. Center inset shows an enlarged view of the 95% confidence intervals for the volume where uncorrected, realigned, and DRIFT have the smallest difference. Right inset shows the regression of framewise displacement to PD. Slopes are shown above as beta values.

At each time point, the realigned data had significantly lower PD compared to the uncorrected data (Average PD_diff_=5.2, p < 10^−5^). Likewise, at every time point, DRIFT had an even lower PD than realignment. The average PD_diff_ between DRIFT and realigned data was PD_diff_ = 0.8, and the volume wise difference was significant at every point with p < 10^−5^. Further, DRIFT performs much more consistently than realignment, as the variance of the PD over all volumes was 0.0008 for DRIFT, and 0.1744 for realignment. The insert to the right of Figure 9 contains a scatter plot of PD and displacement for uncorrected, realigned and Drift corrected volumes, with the slopes indicated as beta values. While displacement is associated strongly with the magnitude of the motion artifact in the uncorrected data, this association weakens by applying realignment to the fMRI data, and is completely removed using DRIFT, verifying the effectiveness of the DRIFT in removing realistic in-plane motion.

### Evaluating DRIFT using a rotating phantom

In our final experiment, we present the results of our rotating phantom inside the scanner as the proof of concept that DRIFT can potentially be used to attenuate motion artifacts during a real fMRI data acquisition. The top row of Figure 10 shows the reconstructed images using the in-house developed reconstruction pipeline for motionless, rotated, realigned, and DRIFT corrected slice, respectfully. The bottom row of Figure 10 plots the mutual information computed between the reference volume and all other volumes acquired throughout the one minute scan for uncorrected (in blue), realigned (in orange) and DRIFT corrected (in green) slices. As demonstrated in this figure, while there is some fluctuation in the computed mutual information for the first 10 slices, the three different reconstructions produced identical mutual information at a mean value equal to 1.82, which provides the maximum possible mutual information for this data, given the level of noise present. Once the phantom starts moving, a higher mutual information indicates better removal of motion artifacts. Throughout the 40 second moving period, realigned images had higher mutual information than uncorrected images in 81% of the timepoints. DRIFT-corrected slices had in the highest mutual information for 100% of the volumes compared to uncorrected data, and in 90% of the volumes compared to realigned data. During the last 10 seconds of motionless scanning, there are differences between the reconstructed, realigned, and DRIFT corrected slices, which is due to the random orientation of the phantom when it stops moving. Finally, the box plot Figure 10 illustrates the significant differences between aggregated mutual information obtained from the uncorrected (in blue), realigned (in orange) and DRIFT-corrected (in green) slices using bar-plots. For all but the initial 10 volumes, realignment significantly increased the mutual information by an average of 0.11 over the uncorrected data (p < 10^−5^). DRIFT-corrected slices also significantly increased the mutual information by an average of 0.15 over the uncorrected data (p < 10^−5^), and 0.04 over the realigned data (p < 10^−5^). Given that the lowest mutual information for any method was 1.25, and the ceiling was 1.82, the total range of the mutual information was only 0.57. The increase in mutual information due to drift DRIFT compared to uncorrected data is 26% of the possible mutual information range, while the increase between DRIFT and realigned data is 7% of the possible range. Together, these results show the superiority of DRIFT in real fMRI scanning to correct for motion contamination during slice acquisition.

**Figure 10:**
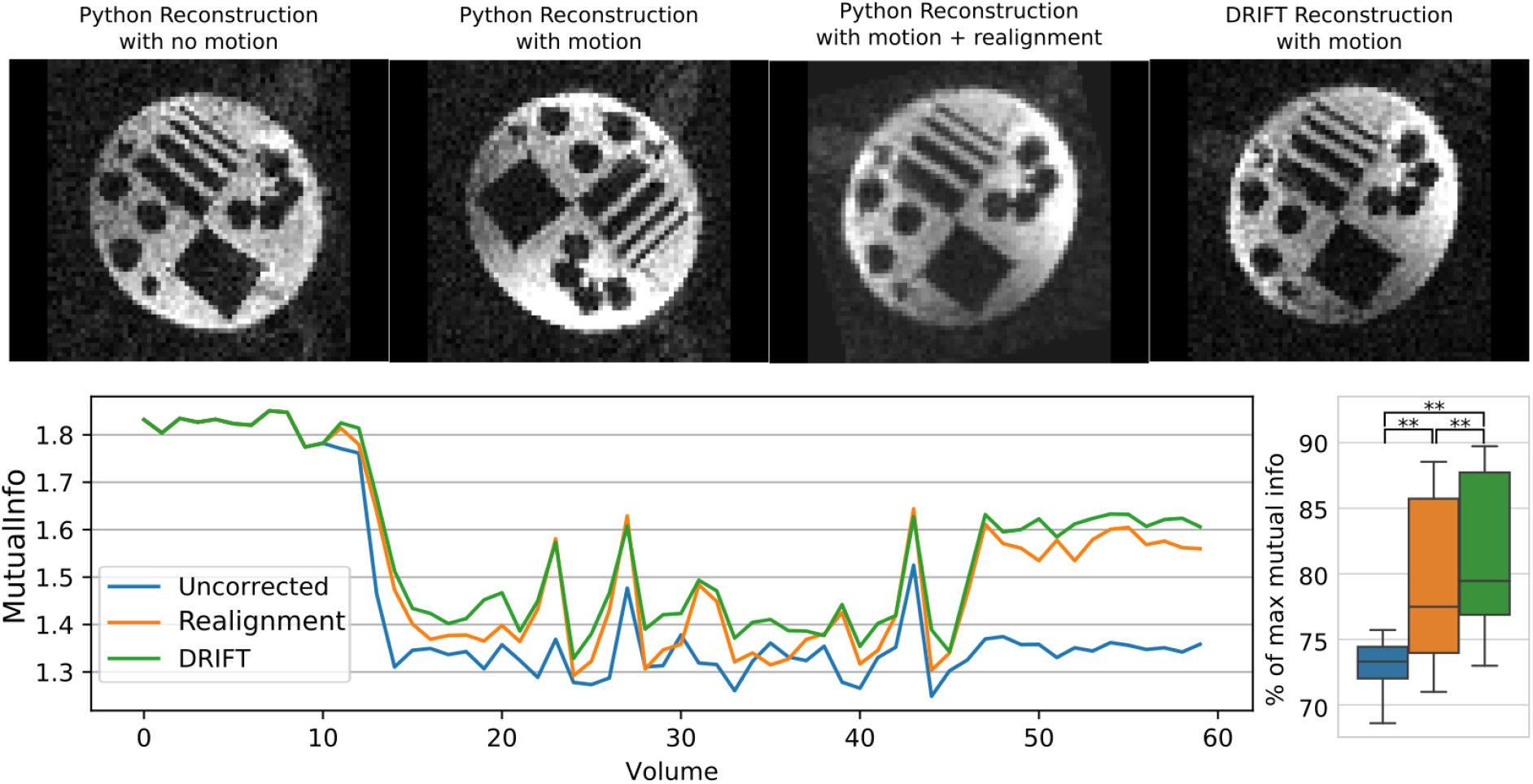
Mutual information of uncorrected, realigned, and DRIFT corrected images in real data. (Top) From left to right: 1: An example of a python reconstructed slice without motion. 2: A rotating slice reconstructed with python. 3: The same rotated slice, corrected with realignment. 4: The same rotated slice, corrected with DRIFT. (Bottom) The mutual information between each volume of real data to the reference volume. The reference volume is created by averaging the first 10 motionless volumes of the scan in the uncorrected dataset.

In this data set, the improvement of drift is hard to see on an individual volume, due to the small nature of the artifact as well as the presence of other noise sources. However, taking a temporal average of all motion contaminated volumes reveals subtle differences between DRIFT corrected volumes and realigned volumes. These averaged volumes are shown in Figure 11. Visually, the DRIFT reconstructed data showed higher contrast and sharper edges, as compared to traditional realignment which generated blurred edges and lower contrast. These “blurred” edges are not an artifact of smoothing, nor are they the result of improper motion estimates – DRIFT and realignment used identical motion parameters for correction. These misalignments are subtle distortions present due to uneven k-space sampling, much like the distortions seen around the edges of simulated data in Figure 7. DRIFT is able to remove many of these artifacts, improving the contrast and reconstructing a more accurate image.

**Figure 11:**
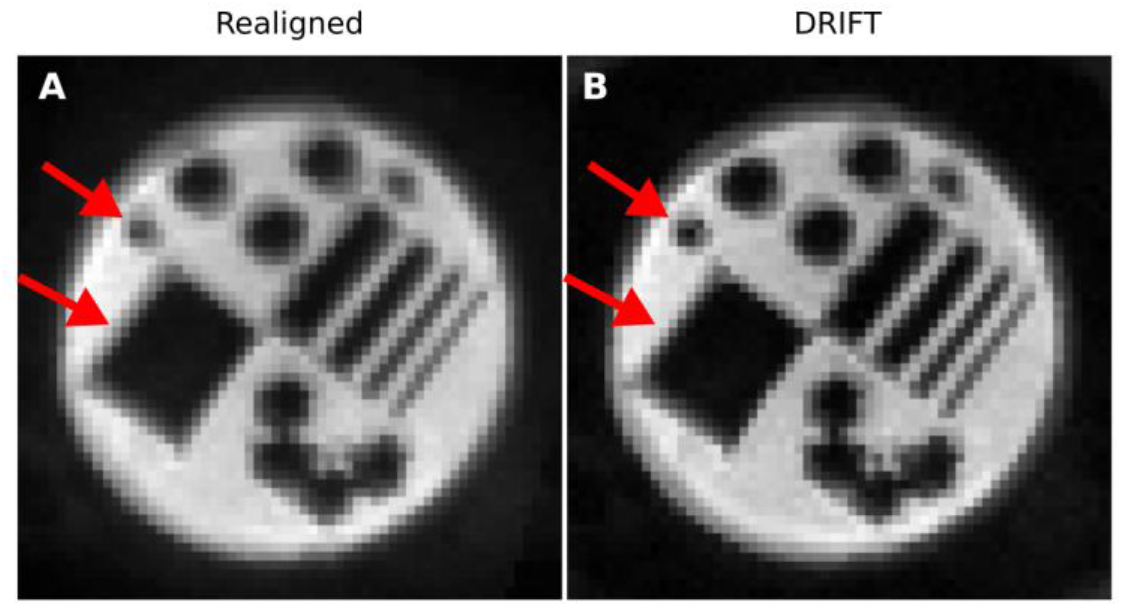
Comparison of time-averaged volumes from corrected scans for realigned data, Drift-corrected data with smoothing, and DRIFT-corrected data without smoothing. The 50 motion contaminated scans from the real data are corrected using either realignment or DRIFT. These volumes are then averaged over time. DRIFT corrected data has noticeably sharper edges. Identical motion parameters were used to correct for motion in DRIFT and realigned data, meaning that the blurring on realigned data is not just due to a poor motion parameter estimate. The sharpness of the DRIFT corrected image is due to the removal of motion artifacts throughout the scan. Removing the smoothing step enhances the

## Discussion

Current strategies that attempt to regress out or remove motion related variance fail to address the motion problem from first principles, and therefor do not successfully model the motion. Motion artifacts are varied and are difficult to describe with just one regressor. Increasing the number of modeled motion effects can alleviate this problem, however this does not address the simple fact that removing motion related variance does not guarantee that the underlying signal is recoverable. Attempting to remove motion artifacts on a reconstructed signal is fundamentally flawed. Motion is not simply an additive noise, as shown through our simulations, and without proper correction, will destroy the underlying signal. By correcting for motion artifacts in the raw data itself, we have the best likelihood of reconstructing an uncorrupted timeseries.

In this paper, we demonstrated the theoretical and practical validity of removing nearly all motion artifacts from fMRI scans. We separately examined common motion found in a typical fMRI scan to investigate reconstruction artifacts which were motion-type specific. We performed a theoretical formulation of the effect of each type of motion using the principles of MR signal generation and the Bloch equations. Next, using the simulated data, we demonstrated the validity of our theoretical findings. Based on our formulation and simulation findings, we argued that neither prospective nor retrospective motion correction methods can completely remove the motion artifacts. Therefore, using a hybrid motion correction method seems to be unavoidable. We then proposed a retrospective method that in combination with the existing slice-based prospective motion correction method, could potentially remove the remaining motion artifacts completely. Again using simulated data we demonstrated clear and substantial improvements in image reconstruction using our proposed retrospective method over other common motion correction routines.

### In-plane motion simulation

For all in-plane motion profiles, DRIFT completely removed all motion artifacts from the image. For in-plane rotation around the *z* axis, the motion artifacts are due to a k-space sampling offset similar to that of Figure 2 (c). Realignment is unable to correct for these artifacts, and only adjusts for any apparent misalignment. Surprisingly, in-plane rotation was not the most destructive form of in-plane motion. Translation along both *x* and *y* generated a larger PD measure than rotation around *z*, even though translation only disrupts the phase of the signal, and preserves the k-space trajectory.

The combination of motion profiles (*y* translation with *z* rotation, and *x* translation with *y* translation) increased the level of motion artifact, but not additively. For example, the PD for translation along *x* during motion was 14.7 for uncorrected data, while the PD for translation along *y* was 13.8. However, the combination of both *x* and *y* translation only increased the PD to 17.0, implying that the errors do not simply add on top of each other, but rather have some form of interaction. This held true for the rotation and translation motion combination as well.

### Out-of-plane motion simulation

For out-of-plane rotation, while it was expected that it would have less error than in-plane rotation, what was surprising was how the out-of-plane rotation artifact was visually nothing like the in-plane rotation artifact. Instead, a rotation around the *y* axis looked like a slight compression of the image along the *x* axis, while a rotation around the *x* axis resembled a slight compression of the image along the *y* axis. This agrees with a qualitative analysis of out-of-plane rotation, which can show that the gradients experienced by the object become compressed spatially during the rotation.

Out-of-plane motion proved to be more destructive in the volumes after motion than the volume during motion. During motion, the uncorrected PD for out-of-plane motion is always lower than the uncorrected PD for in-plane motion. However, this does not mean that in-plane motion is more harmful than out-of-plane motion. In general, in-plane motion has a higher PD in the “during motion” scan, but after motion these artifacts can be optimally removed through PMC, DRIFT, or to a lesser degree, realignment. Conversely, out-of-plane motion has relatively low PD in the “during motion” scan, but has lasting spin-history artifacts in the multiple volumes after motion that cannot easily be corrected for by any method, and so the uncorrupted signal is not recoverable. In the volumes after motion, DRIFT and uncorrected data have a higher PD than realigned data. Much of this is due to misalignment; DRIFT operates on the k-space of acquired slices, and thus cannot correct for out of plane misalignment. However, some of this error is also due to spin-history artifacts, and are responsible for most of the PD in the realigned data. This highlights that spin-history artifacts are an entirely different kind of artifact, unrelated to misalignment or k-space readout artifacts. Because of this, spin-history artifacts are extremely challenging if not impossible to correct for. Given these observations, spin-history artifacts should still be considered as more harmful than in-plane motion artifacts. However, this does strengthen the argument that fMRI needs slice-based PMC to successfully address the motion problem.

Interestingly, realignment increased the PD for the “during motion” volume contaminated with out-of-plane motion. In theory, and depending on the difference measure being evaluated, realignment software should never result in a higher percent difference than uncorrected, as it runs a cost-function minimization routine. If all attempted orientations result in a larger error than the original position, no realignment will be done, and the error should be at most equivalent to uncorrected data. In our reconstructions, the realignment software was given the true motion parameters rather than allowing it to estimate the motion itself. This means that correcting for the true motion does not consistently reduce image differences when motion occurs across slices within a single volume. Because of this, difference minimizing cost functions may provide very unreliable motion estimates for volumes with high motion. This highlights the harmful effects of intra-volume motion and slice misalignment.

Out-of-plane translation and rotation *without* spin-history artifacts proved to be the least destructive of all motion types in our simulations. The simulation of out-of-plane translation is a critical piece of evidence for our motion correction method. It effectively demonstrates that out-of-plane translation only causes artifacts if it causes the following slices to be over-or under-excited. PMC is already demonstrated to be accurate to around 0.3mm with optical and navigator tracking (Dold *et al.*, no date; Speck, Hennig and Zaitsev, 2006; White *et al.*, 2010), and has shown great promise in its ability to compensate for motion on a slice by slice basis (Speck, Hennig and Zaitsev, 2006; Beall and Lowe, 2014). Thus, with proper PMC, no additional motion correction of any kind would be necessary for out-of-plane translation.

### Simulation with real subject motion

For the previous 10 simulated scans used to demonstrate the different types of in-plane and out-of-plane motion artifacts in Figure 7 and Figure 8, the motion occurred entirely over one slice acquisition (two TE’s). The motion we used consisted of either 5mm translations or 5 degree rotations, which occurred along various axes. While it is rare for a subject to move 5 degrees or 5mm within a period of two TE’s, in real fMRI scanning, this level of motion does fall well within physiologically possible ranges. Nonetheless, we were also able to show the superiority of our method on a simulated scan using a real subject’s motion profile. In this particular simulation, not only was the motion taken from a real subject, but also it was simulated to happen gradually over the entire TR, not just during the readout of a single slice. DRIFT was still able to significantly improve the reconstruction in this realistic simulation.

The simulation with the subject’s real motion parameters has smaller translations and rotations than the other 10 simulations, however it has the largest uncorrected error with a peak value of 16.1%, which is higher than any of the exaggerated motion errors. There are multiple explanations for this: first, though the relative volume-to-volume displacements are smaller than 5 degrees or 5mm, over the course of the simulation, the absolute displacement relative to the reference volume accumulates up to a total of 4.1 degrees of rotation and 2.3 mm of total translation at volume 22. Second, we demonstrated in the other simulations that the combination of motions is the more destructive than each motion individually. This simulation has three motion types occurring simultaneously: *z* rotation, *x* translation, and *y* translation, while our simulated data only ever had two motion types occurring simultaneously. Finally, this PD is calculated over the entire volume, rather than just a slice. This increases the number of high-contrast edges that contribute to large PDs.

### Real data correction

For real data, we were able to show the benefit of DRIFT when added to the raw data processing pipeline. While the benefit may seem small, this experiment could be drastically improved to further enhance DRIFT. Primarily, exact sub-TR motion tracking was not available for this scan. If the motion is not accurately measured, this will cause additional errors in DRIFT’s k-space mapping calculations. For this study, we took the relative deflection of the phantom between two volumes and estimated k-space rotations assuming constant rotation. In reality, it is possible the rotation experienced acceleration or deceleration during the TR. Further, DRIFT works best when working directly with the gradient profiles in the exact pulse sequence, as with POSSUM’s simulated data, or at the very least a full k-space trajectory map. Our pipeline estimated a full 128×64 k-space trajectory from a single exemplar read line provided in the raw file header. Without exact knowledge of both the pulse sequence and the motion, we rely on estimating both of these parameters, which have detrimental effects on the results. Finally, the Siemen’s raw data processing pipeline consists of approximately 20 to 30 signal processing steps, which take advantage of the pulse sequence design, adjustment volume for phase corrections, and several other system-dependent inputs that are used for a more accurate reconstruction. Our pipeline consisted of only 3 main processing steps. Integrating DRIFT into the full system pipeline would maximize its effectiveness.

If DRIFT can be integrated directly into a scanner’s system, it can be used with any number of different pulse sequences. One particularly useful area would be 3D EPI. 3D EPI is a technique in which an entire volume is acquired in one RF pulse, and a three dimensional k-space trajectory is traced and sampled. This essentially makes the reconstruction a single 3D FFT rather than multiple 2D FFTs. Using the principles developed in this paper, our reconstruction algorithm could theoretically be applied directly to 3D sampled data, completely eliminating all artifacts during sampling. This would actually eliminate the need for PMC completely, as DRIFT could also be used to re-orient each volume, providing accurate motion estimates were still being recorded. Future work could involve developing this method for a 3D pulse sequence.

## Conclusion

We have developed a novel technique that both corrects for motion while simultaneously reconstructing the data. This method significantly attenuates the motion artifacts on a slice-by-slice basis. By correcting for artifacts in the raw k-space data, we are able to completely remove motion related artifacts and restore the original signal if the steady-state is not disturbed. We demonstrated in both real and simulated data that DRIFT lowers the error between motion contaminated volumes and the reference volume, indicating a better realignment. This was tested and verified over a variety common motion profiles in fMRI. To extend the application beyond theory and simulation, a real fMRI scan was acquired and corrected with our method. DRIFT’s advantage over other motion correction methods is the fact that it corrects for motion in the raw data reconstruction itself. Not only does this eliminate the need for interpolation based motion correction in the reconstructed scan, but also, based on the properties of fMRI reconstruction and the Fourier transform, this is the only way to completely remove motion artifacts from the fMRI signal. DRIFT, when combined with modern prospective motion correction, could potentially eliminate all motion artifacts from fMRI scans.

## Appendix A

## Derivation of the fMRI signal equation without motion

The Bloch equation states that transverse magnetization at a 3D spatial coordinate ***r*** at time *t* (relative to the last RF pulse at *t*_*0*_) is equal to the magnitude of the transverse magnetization at the time of the RF pulse (|*M*_*xy*_(***r***, ***t***_0_)|), multiplied with a phase component described by its rotation *e*^*iω*(***r***,*t*)^, and it’s T_2_* decay described by 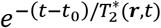, where ω is the Larmor frequency of the protons at point ***r***, and T_2_*(***r***,*t)* is the rate of decay of the transverse magnetization, which varies spatially over ***r*** as well as temporally, which is the basis of the BOLD signal. The full equation is described in eq. 1:

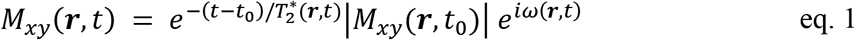

The signal acquired by the receive coils in an fMRI scanner is equal to the sum of all transverse components from all the protons in the excited slice. In other words, we take an integral of this equation over all points in ***r*** for excited tissues. We can simplify this equation if the readout time is small compared to the T_2_* decay by omitting the first exponential. Additionally, once the objects flip angle reaches steady state, Mxy at time t0 is assumed to be constant for each location ***r***, with spatial variation as a function of proton density, ρ. With these adjustments, eq. 1 can be approximated to create a signal equation:

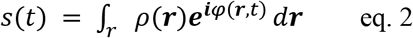

Where *φ* is the accumulated phase, which is derived from the function describing the frequency at which the protons spin in the scanner, *ω*(***r***, *t*):

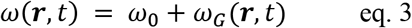

Where *ω*_0_ is the frequency due to the static magnetic field, B0, and *ω*_*G*_(***r***, *t*) is the frequency offset due to any spatial gradients applied in the scanner. The magnitude of this gradient-induced phase offset varies depending on the position ***r*** in the scanner. This is the fundamental principle behind frequency encoding, which allows for image reconstruction in EPI pulse sequences. In a typical scanner, this phase is demodulated, essentially removing the phase component belonging to *ω*_0_, leaving only the *ω*_*G*_ component. The actual amount of frequency offset depends on the gradient strength, the location of the object, and the gyromagnetic ratio, shown in eq. 4:

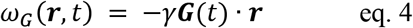

Where the negative sign accounts for the “Counter clockwise positive” rotation convention. A typical EPI pulse sequence takes about 60ms to acquire a single 2D slice of k-space. During this time, the frequency offset accumulates, referred to as “accumulated phase”, which is calculated as the integral of the frequency offset over the readout time:

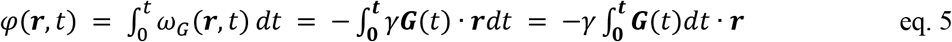

The factors γ and ***r*** do not change with time, and so they can be placed outside the integral. This leads to the classical description of “k-space”, where the signal at time t is being recorded from location ***k*** in k-space:

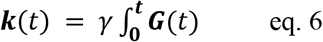

Substituting equations eq. 6 and eq. 5 into eq. 2, we obtain:

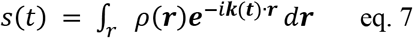

which has the form of a Fourier transform.

## Derivation of the fMRI singal equation with motion

This equation is where we begin our analysis of motion in fMRI. We can now consider two coordinate systems: The object, ***r***_*ob*_, and the scanner, ***r***_*sc*_. Without motion, these coordinate systems are static, and we simply set them equal to each other, where ***r*** = ***r***_*ob*_ = ***r***_*sc*_. However, this no longer holds true if the object moves. When motion is present, we must describe the object’s location relative to its initial position, specifically at the time of excitation from the most recent RF pulse. If the object is initially aligned with the scanner’s coordinate system, ***r***_*sc*_, and moves in a way described by a rotation matrix R and a translation vector ***T***, we can describe a point in the object’s coordinate in terms of the scanner’s coordinates:

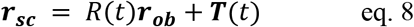

And it follows that we can describe a point in the scanner in terms of the object’s coordinates:

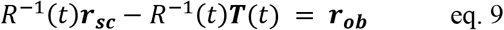

To model the motion as a stationary object with an inverse rotation of the scanner, we must reevaluate eq. 4, eq. 5, and eq. 6 to account for this rotation, as the spatial coordinate ***r*** is now changing with time, and cannot be removed from the integral in eq. 5:

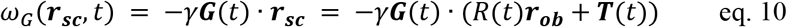

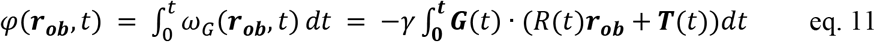

We now distribute the dot product with ***G*** and evaluate each component separately:

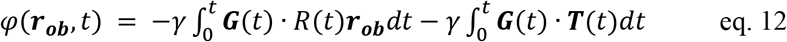

The first term deals with rotation and has the actual object’s coordinates in the integral term. We can use the properties of a dot product to simplify this notation. The geometric definition of a dot product of two vectors in space is

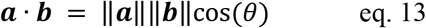

Where ‖***a***‖ is the magnitude of the vector, and theta is the angle between **a** and **b**. In our example, ***r***_*ob*_ is rotated by the rotation matrix R. Because a rotation will not change the magnitude of a vector, the first two components of eq. 13 remain unchanged. The angle between the vectors, however, will change depending on the rotation matrix. A rotation matrix R rotating vector ***b*** by α degrees away from vector ***a*** changes eq. 13 to:

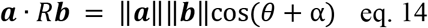

Likewise, rotating vector ***a*** by α degrees away from vector ***b*** also increases the angle between the two vectors by α, and the result is the same as equation eq. 14. Note, that rotating vector ***a*** away from vector b is the inverse (opposite) rotation applied to rotate vector ***b*** away from vector ***a***. For rotation matrices, a rotation in the opposite direction is equivalent to the inverse of the original matrix. In other words:

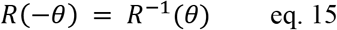

From eq. 14, this means:

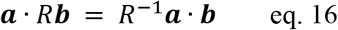

Using this property in eq. 12, we can move the rotation to the gradient vector ***G***, and once again remove the ***r***_*ob*_ vector from the integral:

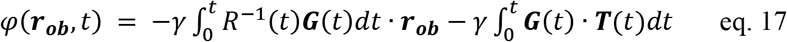

We can now modify our K-space formula and update our signal equation:

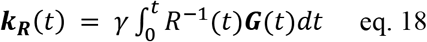

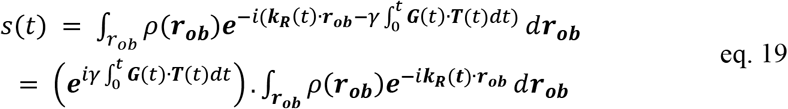

